# Bacterial H-NS contacts DNA at the same irregularly spaced sites in both bridged and hemi-sequestered linear filaments

**DOI:** 10.1101/2020.06.11.146589

**Authors:** Beth A. Shen, Christine M. Hustmyer, Daniel Roston, Michael B. Wolfe, Robert Landick

## Abstract

Gene silencing in bacteria is mediated by chromatin proteins, of which *Escherichia coli* H-NS is a paradigmatic example. H-NS forms nucleoprotein filaments with either one or two DNA duplexes. However, the structures, arrangements of DNA-binding domains (DBDs), and positions of DBD–DNA contacts in linear and bridged filaments are uncertain. To characterize the contacts that silence transcription by RNA polymerase, we combined ·OH footprinting, molecular dynamics, statistical modeling, and DBD mapping using a chemical nuclease (Fe^2+^-EDTA) tethered to the DBDs (TEN-map). We find that H-NS DBDs contact DNA at indistinguishable locations in bridged or linear filaments and that the DBDs vary in orientation and position with ~10-bp average spacing. Our results support a hemi-sequestration model of linear-to-bridged H-NS switching in which linear filaments able to inhibit only transcription initiation switch to bridged filaments able to inhibit both initiation and elongation using the same irregularly spaced DNA contact sites.

**Highlights:**

- Tethered-nuclease mapping (TEN-map) of H-NS DNA-binding domains detects DNA contacts
- Bridged and linear H-NS filaments use the same DNA contact sites
- H-NS–DNA contacts are unevenly spaced with ~10 bp average separation
- AT-steps, minor groove width, and electrostatic potential best predict contact sites

## INTRODUCTION

Extensive study of nucleosome-based chromatin structure has provided broad insight into mechanisms of chromatin-mediated gene regulation in eukaryotes (Cook and Marenduzzo, 2018; Luger et al., 1997; Talbert et al., 2019). For bacteria, however, an analogous understanding of chromosomal gene regulation remains elusive in part due to the diverse and complex DNA-binding properties of the multiple DNA-binding proteins (DBPs) that generate bacterial chromatin (Dame et al., 2020; Grainger, 2016; Shen and Landick, 2019). The γ-proteobacterial <u>h</u>istone-like <u>n</u>ucleoid <u>s</u>tructuring protein (H-NS; 15 kDa monomer) provides a paradigmatic model for this complexity (Grainger, 2016; Qin et al., 2019). H-NS, which contains two oligomerization sites in its N-terminal domain (NTD) and a C-terminal DNA binding domain (DBD), forms gene-silencing, oligomeric filaments on ~15% of the *E. coli* genome by targeting AT-rich DNA, notably on pathogenicity operons (e.g. SPI-1 in *Salmonella*), surface-antigen operons (e.g. LPS biosynthesis in *Escherichia coli*), and horizontally acquired genes (e.g. *bgl* operon in *E. coli*) (Figure 1A,B) (Gawade et al., 2020; Grainger et al., 2006; Kahramanoglou et al., 2011; Lucchini et al., 2006; Navarre et al., 2006; Sankar et al., 2009).

**Figure 1.**
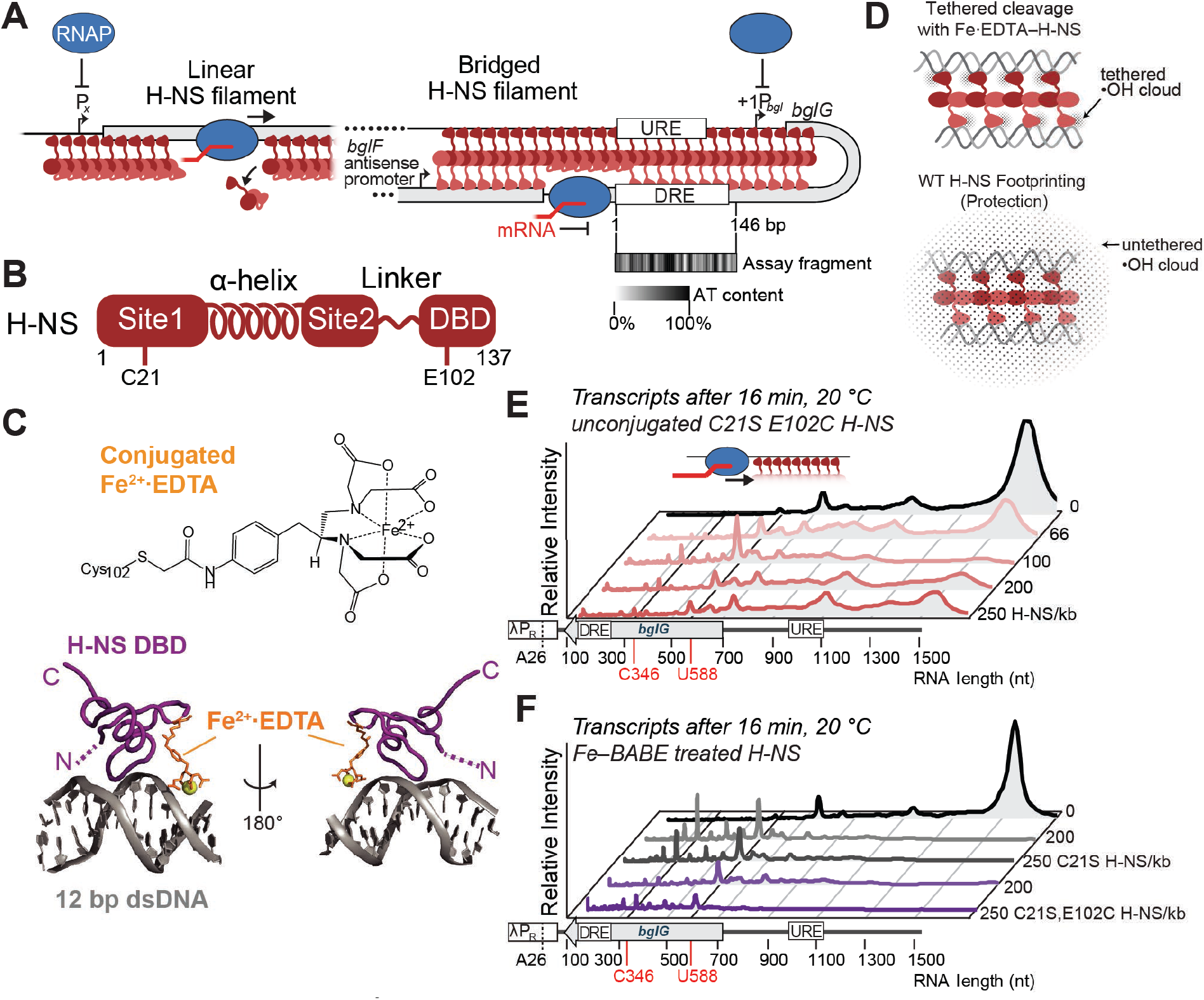
H-NS tethered to Fe^2+^·EDTA forms bridged filaments that stimulate pausing by RNA polymerase. (A) Mechanisms of gene silencing by H-NS illustrated for the *E. coli bgl* operon. Linear and bridged H-NS filaments (dark and light red monomers) can block RNAP (blue) from binding to promoters whereas only bridged H-NS can inhibit elongation of RNAP by stimulating pausing (Dame et al., 2002; Kotlajich et al., 2015; Shin et al., 2005; Singh 2013). AT-rich, high affinitybinding sites nucleate filament formation at the *bgl* upstream and downstream regulatory elements (URE and DRE) (Dole et al., 2004). The AT-content of the 146-bp DRE DNA fragment used in the TEN-map assay is shown using a gray-scale heat map. (B) Structure of the H-NS monomer. Sites 1 and 2 in the N-terminal oligomerization domain, connected by a flexible α-helix, make head-head, tail–tail contacts in H-NS–DNA filaments (Shahul Hameed et al., 2018; van der Valk et al., 2017). A 15-aa flexible linker connects the C-terminal DBDs to the tail–tail junction (Gao et al., 2017). Substitutions (C21S, E102C) enabling Fe^2+^·EDTA conjugation are indicated. (C) Model of the H-NS DBD (PDB 2L93) bound to 12-bp duplex DNA (Riccardi et al., 2019) with Fe^2+^·EDTA tethered to position 102. (D) Comparison of ·OH generated in TEN-map (top) vs. ·OH footprinting (bottom) assays. (E) Pseudo-densitometry traces of RNA transcripts formed by RNAP at 20 °C and 30 μM NTP on a 1.5 kb *bgl* DNA 16 min after release from a λP_R_ promoter–A26 halt site with no H-NS (top) or 66–250 C21S, E102C H-NS monomer/kb DNA. Peaks represent paused RNAP (e.g., at positions 346 and 588) or run-off transcript (1538 nt) (Kotlajich et al., 2015). At ~50 to ~150 monomer/kb, H-NS forms bridged filaments that stimulate pausing whereas higher concentrations of H-NS saturate DBD binding sites in linear filaments (Boudreau et al., 2018; Kotlajich et al., 2015). Bridged and linear filament formation was verified by EMSA (Figure S2). See also Figure S1 for transcription results using wild-type H-NS and Figure S3 for sequence of *bgl* DRE. (F) RNA transcripts formed in the same assay as panel E but using control C21S H-NS and C21S, E102C H-NS treated with FeBABE. Fe^2+^·EDTA–H-NS formed by C21S, E102C H-NS contained Fe^2+^·EDTA conjugated to ~50% of the E102C thiols (see Methods).

H-NS dimers oligomerize on DNA through head–head and tail–tail NTD contacts into filaments that adopt either linear or bridged conformations in which H-NS interacts with one or two segments of duplex DNA, respectively (Figure 1A,B) (Arold et al., 2010; Dame et al., 2006; Dame et al., 2000; Gordon et al., 2011; Kotlajich et al., 2015; Liu et al., 2010; Ulissi et al., 2014). The DBD forms an AT-hook-like structure that binds the narrow minor groove of AT-rich DNA (Gordon et al., 2011), consistent with H-NS filament formation on ~15% of the *E. coli* genome with above-average AT content that may reflect acquisition via horizontal gene transfer (Grainger et al., 2006). A flexible linker connecting the DBD and NTD enables DBD–DNA contacts (Gao et al., 2017; Gulvady et al., 2018). However, in some conditions, a subset of DBDs is proposed to interact with the NTD rather than DNA yielding a hemi-sequestered, linear conformation of H-NS–DNA filaments (Shahul Hameed et al., 2018; van der Valk et al., 2017) (Figure 1A).

Bridged or linear H-NS filaments may inhibit transcription initiation by RNA polymerase (RNAP) by occluding promoters (Srinivasan et al., 2013) and bridged filaments may trap RNAP at promoters (Dame et al., 2002; Shin et al., 2005). However, only bridged but not linear filaments inhibit transcript elongation by RNAP, which is proposed to occur via a topological trapping mechanism (Figures 1A and S1) (Boudreau et al., 2018; Kotlajich et al., 2015). Switching between linear and bridged states, DNA avidity, and thus H-NS effects on gene expression may be modulated in vivo by changes in H-NS levels, H-NS filament modifiers like Hha, cellular solutes and thus environmental conditions, and temperature (Boudreau et al., 2018; Kotlajich et al., 2015; Qin et al., 2020; van der Valk et al., 2017; Zhao et al., 2021a; Zhao et al., 2021b).

It remains unclear how the DBDs are arranged on DNA within either linear or bridged H-NS filaments and if the contacts change in the switch between linear and bridged conformations that differentially affect steps in transcription. DNase and ·OH footprinting experiments reveal nucleation at AT-rich high-affinity sites and H-NS oligomerization with an apparent periodicity of DBD contacts every ~10 bp (Bouffartigues et al., 2007; Ulissi et al., 2014; Will et al., 2015). Additionally, single-molecule pulling experiments show that strong DBD–DNA contacts at ~10- or ~20-bp intervals are common in bridged filaments (Dame et al., 2006). A prevalent model for H-NS filament organization based on an NTD crystal structure predicts ~30-bp spacing of DBDs in a bridged filament (Arold et al., 2010). Although H-NS is known to bind AT-rich DNA (Gordon et al., 2011) the locations of the H-NS DBDs in filaments relative to DNA sequence features is not well defined. Because a flexible linker connects the DBDs to the filament-forming NTD, the extent to which DBD–DNA contacts within a filament are regularly spaced is also unclear. New approaches are needed to determine how DNA sequence governs DBD location and the arrangement of DBDs in an H-NS filament.

To gain insight into the organization of DBD locations in filaments, we developed and applied a multimodal approach that combines a novel DBD-mapping method using a <u>te</u>thered chemical <u>n</u>uclease (TEN-map), hydroxyl radical (·OH) footprinting, computational molecular dynamic (MD) simulations, and rigorous statistical analysis and modeling. Our results reveal that DBDs contact DNA irregularly in an H-NS filament but at the same locations in bridged and linear filaments. They also confirm the hemi-sequestration model of linear-to-bridged H-NS switching and providing a conceptual framework for understanding the mechanism of gene silencing by H-NS and related proteins in bacteria.

## RESULTS

### Tethered-nuclease mapping (TEN-map) reveals H-NS DBD locations in H-NS filaments

To determine the spacing and orientation of DBDs within an H-NS filament, we sought a method that could reveal DBD location and orientation on DNA as a complement to conventional ·OH footprinting, which defines regions of protection on DNA but not which part of a protein is responsible for the protection. To that end, we attached the chemical nuclease Fe^2+^·EDTA to the H-NS DBD to enable <u>te</u>thered <u>n</u>uclease <u>map</u>ping of DBD contacts on DNA (TEN-map; Figure 1C,D). TEN-map reveals the locations and orientations of DBDs by generating hydroxyl radicals (·OH) that can cleave the nearby DNA backbone within ~10 Å from the Fe moiety (Cheal et al., 2009; Greiner et al., 1997; Rana and Meares, 1991). For example, this method detects DNA contacts by the *E. coli* RNAP a subunit CTDs that also bind the minor groove of promoter DNA and are tethered by flexible linkers (Murakami et al., 1997).

For TEN-map, Fe^2+^·EDTA was attached to Cys substituted for E102 in the H-NS DBD by reaction with FeBABE (Fe(III) (*S*)-1-(*p*-bromoacetamidobenzyl)ethylenediaminetetraacetate) (Greiner et al., 1997; Rana and Meares, 1991). We chose E102 because it is near DNA but not part of the DBD–DNA interface in a model of the H-NS DBD (Gordon et al., 2011), is predicted to remain solvent exposed in H-NS filaments, and can be replaced by Ala without effect on H-NS DBD binding to DNA (Gordon et al., 2011) (Figure 1C). Modeling of Fe^2+^·EDTA attached to C102 DBD predicted that the chelated Fe^2+^ remains near the DNA but would not interfere with the H-NS–DNA interface (Figure 1C; see below). *E. coli* H-NS naturally contains Cys at position 21 in the NTD site 1 (Figure 1B). Therefore, we constructed, overexpressed, and purified recombinant H-NS C21S E102C. Conjugation with FeBABE yielded the tethered nuclease H-NS (Fe^2+^·EDTA–H-NS).

We first tested whether Fe^2+^·EDTA–H-NS could form H-NS filaments and inhibit transcript elongation by RNAP as observed previously for wild-type H-NS (Boudreau et al., 2018; Kotlajich et al., 2015). We used a 1.5 kb duplex DNA in which the upstream and downstream regulatory elements (URE and DRE) from the H-NS silenced *bgl* operon (Dole et al., 2004; Nagarajavel et al., 2007) were transcribed from a strong λP_R_ promoter positioned to mimic natural transcription of the *bglF* antisense promoter (Figures 1A and 1E). Unconjugated C21S E102C H-NS at ~66 H-NS/kb DNA, conditions found previously to favor bridged filaments, stimulated pausing by RNAP whereas linear filaments formed at ≥200 H-NS/kb had much less effect (Figure 1E; compare to WT H-NS in Figure S1). Both C21S H-NS and C21S, E102C H-NS treated with FeBABE stimulated pausing similarly to wild-type bridged H-NS filaments but required higher concentrations to show effects (Figure 1F). However, even the highest levels of the FeBABE-treated H-NS achievable in this assay (250 H-NS/kb) still inhibited elongation. We concluded that bridged Fe^2+^·EDTA–H-NS filaments stimulate pausing similarly to wild-type H-NS but that the conjugated H-NS either was constitutively bridged or was partially inactivated by the conjugation reaction so that concentrations of active H-NS needed to form linear filaments could not be achieved. Shahul Hameed et al. (2018) report E102 may interact with the H-NS NTD in linear filaments, which could explain constitutive bridging by Fe^2+^·EDTA–H-NS.

To probe the structure of nucleoprotein filaments formed by Fe^2+^·EDTA–H-NS, we visualized them using an electrophoretic mobility shift assay (EMSA) (Figure S2). Wild-type and C21S H-NS formed the slower and faster migrating bands at 100 and 250 H-NS/kb DNA, characteristic of bridged and linear filaments, respectively (Boudreau et al., 2018; Kotlajich et al., 2015). However, neither C21S E102C H-NS nor Fe^2+^·EDTA-conjugated H-NS appeared able to form linear filaments; both exhibited shifts to more slowly migrating filaments at 250 H-NS/kb rather than to faster migrating linear filaments. Both C21S and C21S–E102C H-NS treated with FeBABE required higher concentrations to form filaments, which may reflect partial loss H-NS activity during biochemical manipulation. These results confirm that Fe^2+^·EDTA-H-NS forms bridged H-NS filaments but suggest that these filaments are constitutively bridged. Since even unconjugated C21S–E102C H-NS was unable to form linear filaments, our results are consistent with the idea that E102 contributes to hemi-sequestration of the DBD via interactions with the NTD in linear H-NS–DNA nucleoprotein filaments (Shahul Hameed et al., 2018).

To probe the locations of the bridged H-NS DBDs on DNA with TEN-map, we chose a 146-bp fragment of the *bgl* operon encompassing the DRE that is known to bind H-NS well in vitro (Figures 1A and S3) (Dole et al., 2004; Nagarajavel et al., 2007). The 146-bp DRE fragment could accommodate 10–15 H-NS DBDs (assuming a binding-site size of 10–15 bp) and enabled single-nt resolution of cleavage when analyzed by denaturing PAGE (Figure 2). We probed TEN-map cleavage on both strands of the 146-bp DRE fragment separately by positioning a 5’-^32^P label at one end of the top or bottom DNA strands. After formation of filaments, phosphodiester backbone cleavage *via* hydrogen abstraction from DNA sugars was triggered by addition of H_2_O_2_ and ascorbic acid to generate ·OH by the Fenton reaction (Balasubramanian et al., 1998). We observed robust cleavage patterns for both DNA strands at Fe^2+^·EDTA-H-NS ≥1 μM, confirming specific positioning of the DBDs bound to the DRE fragment (Figure 2A,B). To quantify the cleavage patterns, we calculated peak areas for cleavage signals at each bp by densitometric analysis (Das et al., 2005). Cleavage maxima were spaced at irregular intervals on both strands and remained unchanged in location up to 5 μM Fe^2+^·EDTA–H-NS (Figure 2C,D).

**Figure 2.**
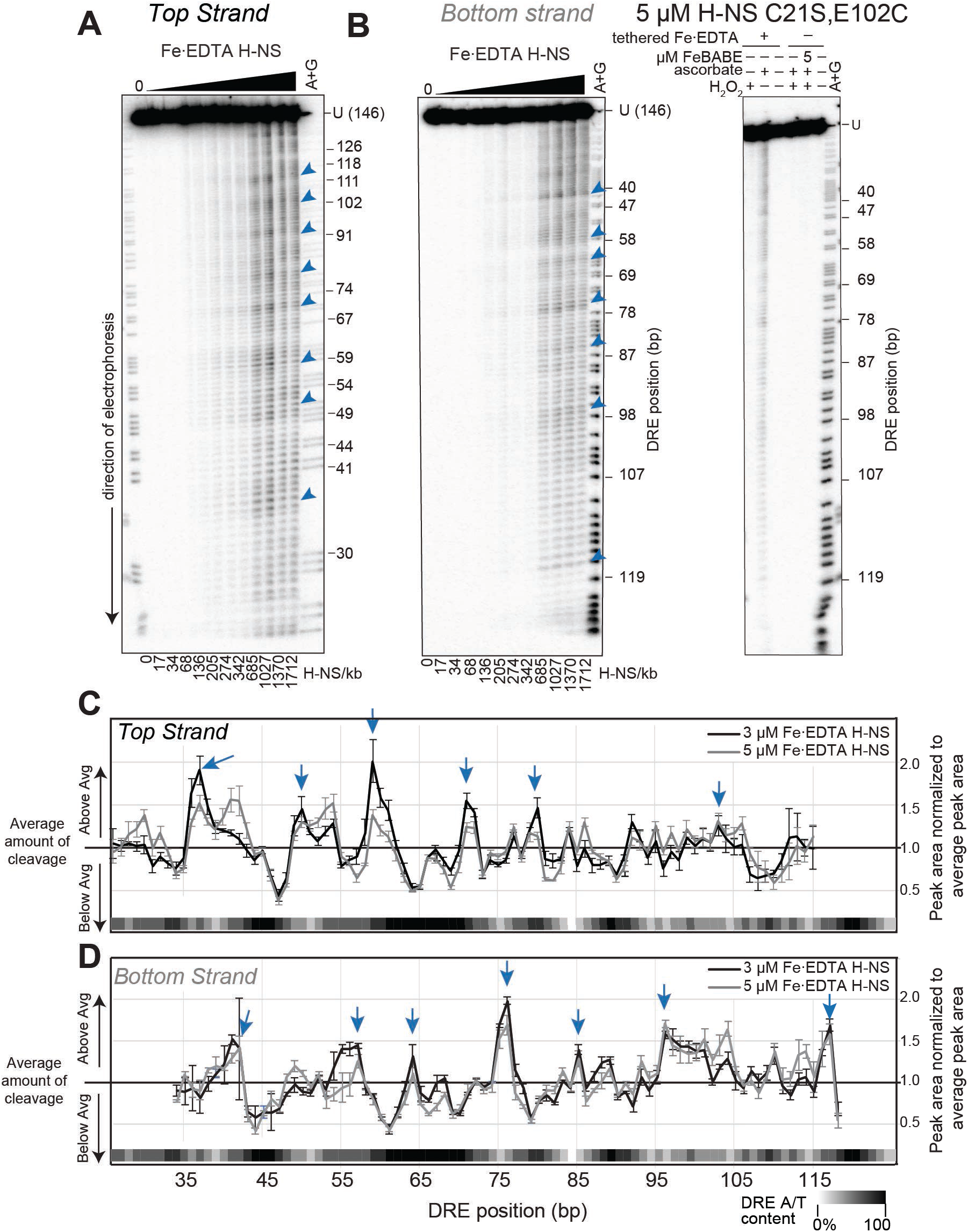
TEN-map locates H-NS DBDs at irregularly spaced sites in the *bgl* DRE. (A) Denaturing 7% polyacrylamide gel showing Fe^2+^·EDTA–H-NS cleavage pattern of the 146-bp DRE DNA labeled on the top strand. Increasing concentrations of Fe^2+^·EDTA–H-NS (50 nM to 5 μM or 17-1712 H-NS/kb) were added to ~20 nM DNA to form filaments. Cleavage was initiated with addition of H_2_O_2_ and ascorbic acid to 2 mM each. Sites of intense cleavage are indicated with blue arrows. A+G, Maxam-Gilbert cleavages after A and G. (B) (Left) Same as panel A, but for the bottom strand. (Right) Control reactions on bottom strand electrophoresed next to cleaved DNA. Little to no cleavage was observed unless all components of the TEN-map assay were present. (C&D) Peak areas calculated by densitometric semi-automated footprinting analysis (SAFA (Das et al., 2005)) and normalized to average signal in each lane for the top (C) and bottom (D) strands (see Methods), shown here for the 3 and 5 μM Fe^2+^·EDTA–H-NS samples. Peak areas above 1 indicate cleavage hot spots (arrows). Error bars show standard deviation of at least three replicates. The AT content of the DRE fragment (5-bp windows) is indicated by the white-black gradient along the X-axis. See Figure S4 for additional TEN-map controls.

To confirm the TEN-map cleavages resulted from Fe^2+^·EDTA tethered to H-NS, we verified that the TEN-map pattern was distinct from ·OH cleavage of free DNA or DNA bound by H-NS when Fe^2+^·EDTA was free in solution rather than tethered (Figure S4). ·OH cleavage using untethered Fe^2+^·EDTA corresponds to conventional ·OH footprint of H-NS and resembled but was not identical to the TEN-map pattern. The resemblance is expected because the bound H-NS DBD excludes ·OH from DBD–DNA interfaces regardless of the source of ·OH; the differences in two patterns confirmed that TEN-map gives information about DBD positioning on DNA based on Fe^2+^·EDTA tethering to one side of the DBD bound in a DNA minor groove; Figure 1C).

Because the DRE fragment used for TEN-map was a short, linear DNA, we asked if either the DNA ends or the DNA fragment length might influence phasing of DBDs along the DNA. We first compared TEN-map cleavage of the 146-bp DRE fragment to that of a 151-bp fragment identical to the 146-bp fragment except for 5 bp added to the ^32^P-labeled end (Figure S5A). We observed identical cleavage patterns for these two fragments, establishing that the DBD locations were determined by the underlying DNA sequence rather than being phased from a nearby DNA end (Figure S5B). We next compared TEN-map of the 146-bp DRE fragment to that of a 549-bp fragment with the same ^32^P-labeled end (top strand) but extended 403 bp in the direction opposite the ^32^P-labeled end (Figure S5C). The TEN-map patterns were essentially identical for these DNAs (Figure S5D), except for a stronger peak at position ~112 on the long DRE DNA fragment. Cleavage at this position may be stronger on the 549-bp DNA because the H-NS filament is stabilized at the position by the adjacent DNA absent on the 146-bp DNA. We conclude that H-NS–DBD contacts are determined by the underlying DNA sequence and not binding of H-NS to flanking regions or to a DNA end.

### H-NS DBD locations are unchanged in obligately bridged filaments or by Mg^2+^

We next sought to verify that the H-NS filaments modified at E102 were always bridged in the TEN-map assay. For this purpose, we first used a previously developed bridging–pulldown assay (van der Valk et al., 2017) to generate unambiguously bridged filaments (Figure 3A and S6). In the bridging–pulldown assay, H-NS–DNA filaments are captured from solution by DNA-coated magnetic beads. Capture requires H-NS bridging between the bead-bound and unbound DNAs. We incubated streptavidin beads pre-loaded with a 685-bp, biotinated bait DNA with a mixture of Fe^2+^·EDTA-H-NS, unmodified H-NS, and unbiotinated 549-bp DRE-containing fragment 5’ ^32^P–end labeled on the top strand (Figure 3A). Bridged filaments, whose formation was unaffected by the presence of conjugated H-NS, were magnetically separated from unattached filaments and TEN-map cleavage was activated (Figure S6A). These obligately bridged cleavage patterns were indistinguishable from those observed without bead-based capture (Figure 3A and S7), confirming that the TEN-map assay reports the locations of bridged H-NS DBDs. As an additional test, we asked how elevated Mg^2+^, which stimulates bridging at concentrations >5 mM (Liu et al., 2010), would affect the TEN-map assigned DBD locations. We found that DBD locations were unaffected by elevated Mg^2+^ (8 mM vs. ~0.3 μM) at either high or low H-NS/DNA ratios (Figures 3B and S8). In agreement with the EMSA assay (Figure S2), these results indicate that H-NS filaments with Fe^2+^·EDTA attached to C102 in the DBD are constitutively bridged.

**Figure 3.**
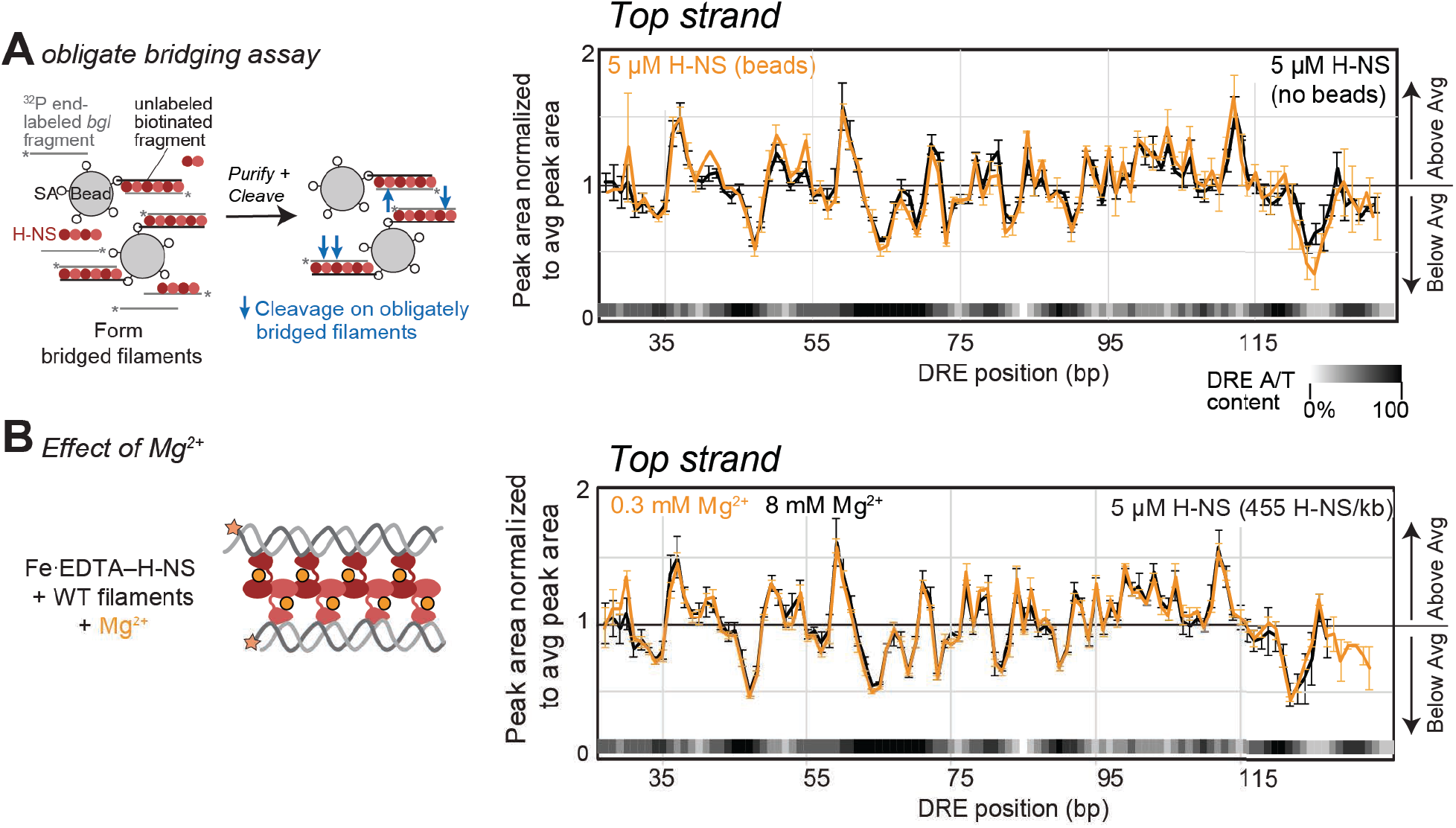
TEN-map reports DBD positions in obligately bridged H-NS filaments. (A) TEN-map cleavage of DRE DNA is unaffected by obligate bridging. Bridged filaments were formed by mixing unlabeled 685-bp DNA fragment bound to paramagnetic beads, 549-bp DRE-containing DNA labeled on the top strand, and H-NS (60% wild-type H-NS + 40% Fe^2+^·EDTA–H-NS) at the indicated concentrations. Obligately bridged filaments were recovered by magnetic separation and resuspended in solution containing H_2_O_2_ to initiate cleavage of DNA (see also Figure S6 and Methods). TEN-map cleavage of the top strand of the DRE region by 5 μM total H-NS (orange) is shown normalized to average peak area in each condition (see Methods). The cleavage pattern is identical to that of the same DNA not tethered to beads (black). (B) TEN-map cleavage of DRE DNA is unaffected by Mg^2+^ concentration. TEN-map cleavage of DRE top strand by Fe^2+^·EDTA–H-NS at 5 μM total H-NS (455 H-NS/kb) in the presence of 8 mM (black) or 0.3 mM (orange) Mg^2+^ and analyzed as described for panel A. See also Figures S6-8.

### Wild-type H-NS DBD ·OH footprints on DNA are unchanged during bridged-to-linear switching

Although the TEN-map cleavage pattern revealed H-NS binding positions within a bridged filament, it could not probe whether the DBDs change locations in the switch from bridged to linear because the derivatized H-NS was constitutively bridged. However, this switch can be probed simply by increasing the concentration of H-NS from ~66 H-NS/kb DNA (bridged filaments) to ≥200 H-NS/kb DNA (linear filaments) (Kotlajich et al., 2015) (Figure 4A). Since the DBDs confer patterns of protection of DNA in conventional ·OH footprinting experiments, we reasoned that assaying ·OH footprints as a function of H-NS concentration should reveal any changes in DBD locations that occur in the switch from bridged to linear filaments. Therefore, we generated ·OH footprints of wild-type H-NS on the DRE encompassing the appropriate ranges of H-NS concentrations (Figures 4 and S9). We normalized ·OH cleavage as a function of H-NS concentration to the unprotected cleavage pattern to correct for intrinsic DNA reactivity (Burkhoff and Tullis, 1988). Strikingly, the H-NS footprint positions did not change as H-NS concentration was increased from a range that favors bridging to well above the concentration that switches filaments to the linear conformation (Figure 4B,C).

**Figure 4.**
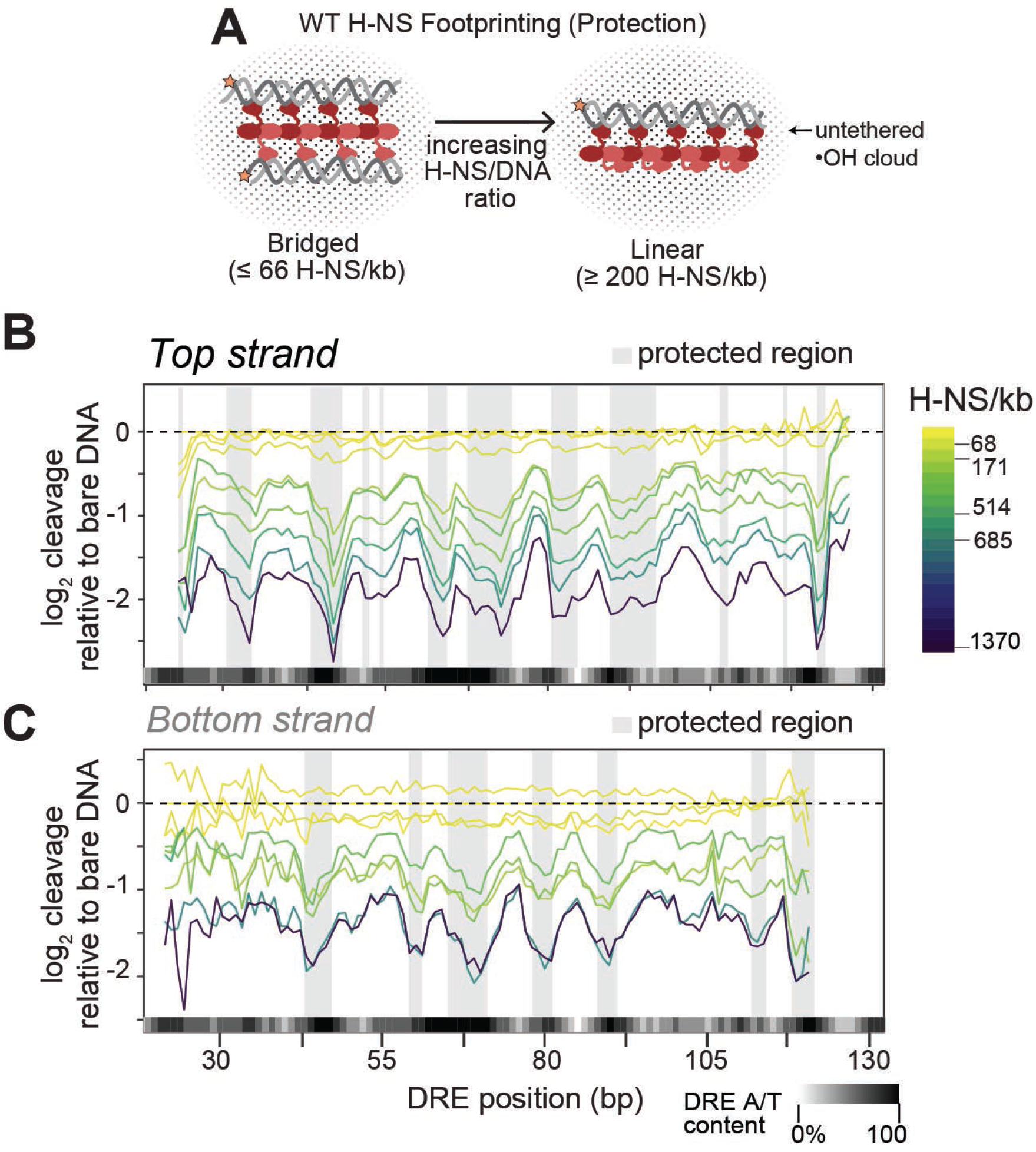
H-NS ·OH footprinting reveals identical protection of the DRE DNA in bridged and linear filaments. (A) Schematic showing footprinting of bridged and linear H-NS filaments generated by increasing H-NS/DNA ratio using ·OH generated in solution (black dots) (see also Figure 1D). WT H-NS switches from bridged to linear filament conformation when H-NS/kb DNA ≥ −200 (Kotlajich et al., 2015) (see Figure S2). (B) Protection patterns on DRE (top strand) at increasing H-NS concentrations (17, 34, 68, 171, 251, 342, 514, 685, 1370 H-NS/kb DNA) are plotted as log_2_ ratios relative to intrinsic cleavage of unbound DNA. Strongly-protected regions are highlighted by shaded boxes (see also Figures S9 and S11). (C) Same as B, but for the bottom strand. In (B&C), lines represent the median of 4 experiments, but error bars are removed for clarity.

As a further test of the conclusion that DBD contacts remain the same in bridged and linear H-NS filaments, we examined previously reported H-NS footprints on different DNA sequences for which H-NS/DNA ratios were reported. Four studies met these criteria (Lucht et al., 1994; Moreira et al., 2011; Ulissi et al., 2014; Will et al., 2018). Despite the use of different footprinting methods (DNaseI or ·OH) and different DNA sequences, no change in apparent DNA contacts at different DNA concentrations encompassing both bridged and linear conditions was evident in any of these studies (Table S1). These prior results thus strongly support our conclusion that bridged and linear H-NS–DNA filaments use the same DNA contact sites with unchanged spacing in the switch from bridged to linear conformations.

### MD simulation and statistical modeling reveal H-NS DBD spacing and orientation

Although both TEN-map and ·OH footprinting indicate unchanged DNA contact sites and spacing in bridged and linear filaments, neither the precise placement nor the orientation of the H-NS DBDs are immediately apparent from the cleavage patterns. Thus, to assign locations and orientations of the DBDs from our TEN-map and ·OH footprinting data, we used a combination of molecular dynamic and statistical modeling. We first generated predicted cleavage and protection patterns for a bound DBD in TEN-map and ·OH footprinting by molecular modeling. For the TEN-map predictions, we modified a structural model of an H-NS DBD bound to the minor groove of DNA (Gordon et al., 2011) to include Fe^2+^·EDTA tethered to the E102C substitution and used it to seed an MD simulation of ·OH generation (Figure 5A). The simulations generated a probability distribution of distances between all possible conformations of the tethered Fe^2+^ and each nucleotide in the duplex DNA strands to which the DBD is bound (arbitrarily designated strands A and B in the model). We assumed that the cleavage propensity at each nucleotide in the DNA fragment is directly proportional to the local concentration of ·OH at that nucleotide, which we estimated by a diffusion model for ·OH centered at the Fe^2+^. Because some nucleotides are shielded from the solvent by the bound DBD, we scaled the concentration of radicals according to the solvent-accessible surface area of the putative cleavage site (Figure 5B) (Balasubramanian et al., 1998). Modeling based on these assumptions yielded an estimate of the relative cleavage propensity at each nucleotide in the duplex DNA bound by the DBD (Figure 5A; see Methods).

**Figure 5.**
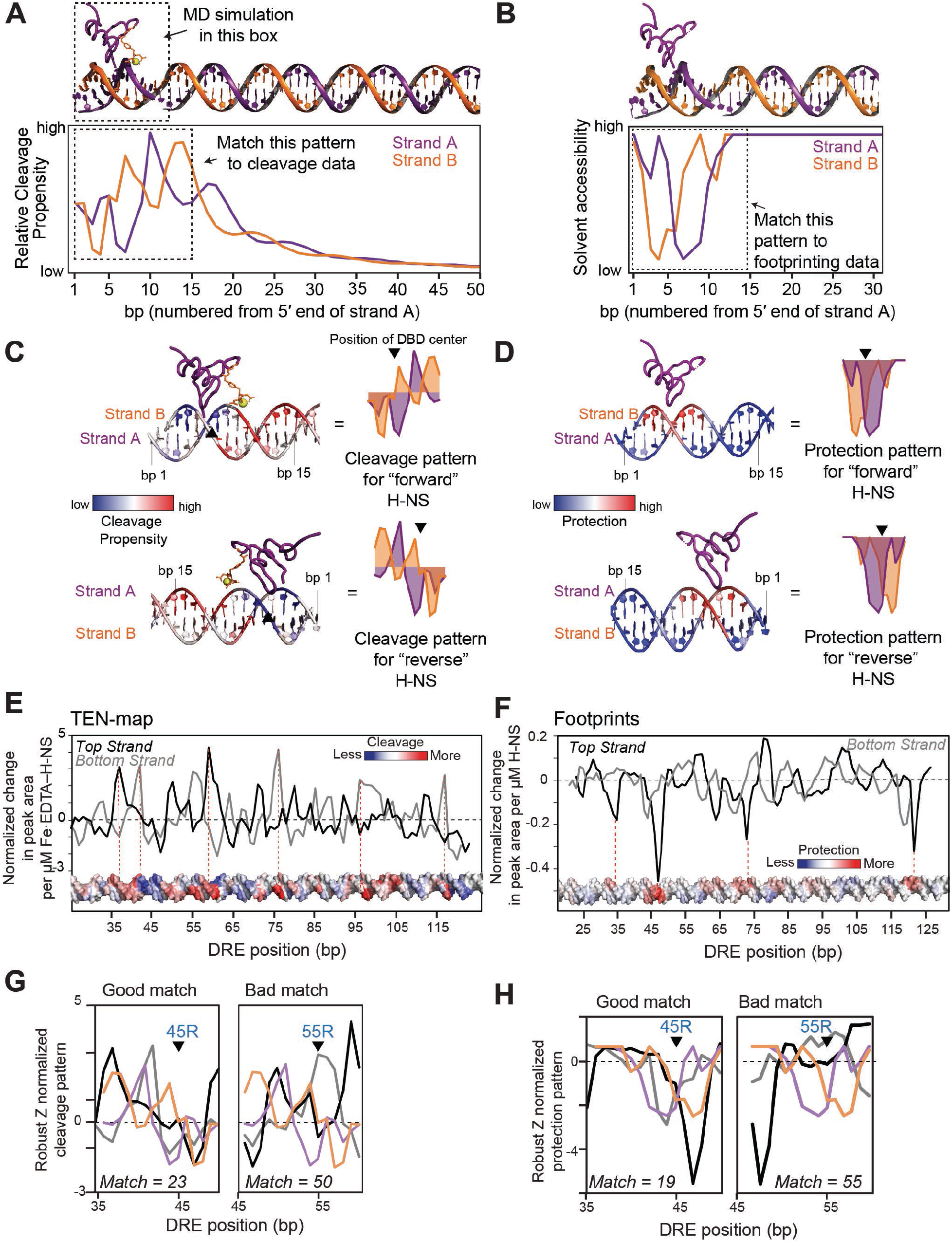
Comparison of predictions from molecular modeling to TEN-map and ·OH footprinting data to place H-NS DBDs on the DRE DNA. (A) MD simulation of Fe^2+^·EDTA (orange) conjugated to the H-NS DBD (purple) bound to a 12-bp DNA duplex (strand A, purple; strand B, orange). The distance between the Fe^2+^ and each bp during the simulation was used to estimate the relative cleavage propensity at each position. Additional DNA (bp 13-50) was included after the MD simulations to estimate cleavage propensities at bp further from the binding site. The cleavage propensity pattern in the dashed box was used in the comparison to experimental TEN-map data. (B) Solvent accessibility of DNA bound by one H-NS DBD calculated from the structure shown at the top of the panel. A 15-bp portion (dashed box) of this pattern was compared to the experimentally determined ·OH footprinting pattern. (C) Overlay of the TEN-map cleavage propensity on B-form DNA with H-NS bound in a forward orientation (PDB 2L93) (Gordon et al., 2011). Cleavage propensity is on a blue-white-red spectrum where blue is low cleavage propensity and red is high cleavage propensity. Cleavage pattern of H-NS bound in the reverse orientation is shown below. The black triangle indicates the center of the H-NS binding site used to align predictions and the cleavage data (see panel G). (D) same as panel C, but displaying the predicted solvent accessibility, or protection, of B-form DNA bound by a single H-NS DBD. Bases more or less protected by H-NS are shown in red or blue, respectively. Slight asymmetry in the protection pattern distinguishes the forward (top) or reverse (bottom) binding directions. (E) Change in TEN-map signal as a function of H-NS concentration (see Methods and Figure S10). More positive slope corresponds to greater H-NS-dependent cleavage. Error in the calculated slopes is in Figure S10. The slope changes are displayed on B-form DNA with red and blue corresponding to greater or lesser extents of cleavage. (F) Change in ·OH footprint signal as a function of H-NS concentration (see Methods and Figure S11). More negative slope corresponds to greater H-NS-dependent protection. The slope changes are displayed on B-form DNA with red and blue corresponding to greater or lesser protection. (G) Examples of alignment of the predicted reverse TEN-map cleavage pattern (panel C) with the TEN-map slope values (panel E) at positions 45 (good match score, 23) or 55 (poor match score, 50). Predicted strand-A cleavage (orange) should match the top strand cleavage pattern (black); predicted strand-B cleavage (purple) should match the bottom strand cleavage pattern (gray). Match scores for the entire DRE fragment are shown in Figure S12. (H) Same as for G but showing alignment of the predicted protection pattern (panel D) with ·OH footprint slopes (panel F).

When the predicted pattern of ·OH cleavage was displayed on B-form DNA, foci of cleavage were evident on each DNA strand offset from the location of the DBD in the minor groove (Figure 5C). The maximum peak of cleavage was predicted to occur at nucleotide +10 of strand A and was flanked by two smaller predicted cleavage peaks in strand B. Adjacent to these sites of high predicted cleavage were sites of protection from cleavage, which resulted from shielding of DNA by the bound DBD. We note that this model does not account for possible contributions to DNA cleavage by ·OH generated from any Fe^2+^·EDTA-DBDs not bound to DNA. These contributions are assumed to be non-specific, to contribute to background DNA cleavage, and thus to be captured in the model by solvent shielding by the bound DBD. In other words, bound DBDs are expected to generate local cleavage patterns that reflect localization of Fe^2+^·EDTA near DNA whereas unbound DBDs are assumed to generate background cleavage from unlocalized ·OH.

We applied a similar analysis to the untethered ·OH footprinting data by using only the calculated solvent exclusion from DNA by the bound DBD to predict protection patterns (Figure 5B). These predicted protection patterns were not completely symmetric due to asymmetric DBD shape when bound to the DNA minor groove in the model (Figure 5D). Thus, although much less robust than for TEN-map, the predicted protection patterns could be compared to ·OH footprints for DBDs bound in either orientation on the DRE fragment.

To generate robust TEN-map and ·OH footprint signals to compare to the predictions, we first used Bayesian linear regression of the DNA cleavage signals as a function of H-NS concentration to assign a single signal strength for each DNA nucleotide on each strand (Figure S10). This analysis was possible because only the intensities and not the locations of the TEN-map cleavages changed as a function H-NS concentration. For the TEN-map data, we centered the normalized cleavage signals for each H-NS concentration at zero (Figure S10A,D), and then fit a Bayesian linear regression model to the signal at each position (Figure S10B). We then used the slopes of these linear-regression fits to generate single-value traces of TEN-map cleavage on each DNA strand for which robust cleavage had a large, positive value (Figures S10C and S10E). Visualization of the H-NS-dependent cleavage on B-form DNA revealed a pattern of cleavage maxima and minima consistent with potential DBD locations (Figure 5E). We used a similar procedure to generate single-value traces of ·OH footprint data on each DNA strand (Figures 5F and S11), for which robust protection had a large, negative value (Figures S11C and S11E).

These single-value traces for both the TEN-map and ·OH footprint results were then compared to the predicted patterns for the respective methods, scanning each nucleotide position on each DRE DNA strand (Figure 5G,H). We calculated Manhattan distances between these predicted cleavage patterns for the DBDs and the observed cleavage traces generated by the linear regression models (Figures S12). The Manhattan distance is the sum of absolute differences between of sets of points; a lower score indicates a closer match (see Methods). The TEN-map and ·OH footprinting match scores revealed similar patterns of DBD interactions with the DRE DNA (Figure S12A and B). To estimate the statistical significance of the match scores, we compared them to values obtained from a Monte Carlo permutation test (see Methods). The match scores could then be converted to *p*-value estimates of the probability they could arise by chance (Figure S12C and D).

To generate a final model of the H-NS filament on the DRE DNA, we combined the *p*-values for the TEN-map and ·OH footprint methods into a single prediction of DBD locations (Figure 6A and S12E). This integrated probability calculation predicted seven DBD locations in the reverse orientation with *p* ≤0.05 and variable spacing of 8-17 bp between the center of the DBD contacts (45R, 62R, 71R, 81R, 93R, 106R, and 114R; 106R was included because its *p*-value was essentially indistinguishable from 0.05). These seven could represent contacts made by a single bridged or linear H-NS–DNA filament (Figure 6B). Three weaker potential DBD contacts in the forward orientation also were predicted (50F, 66F, 88F). The forward-oriented DBDs were offset by ~5 bp from nearby stronger predictions for DBDs in the reverse orientation. This offset prevents steric clash with the more robustly predicted reverse DBDs if the forward DBDs are placed on the DRE DNA (Figure 6B). It is thus unclear whether they might represent alternative contacts of the same DBD or weak contacts by an adjacent DBD.

**Figure 6.**
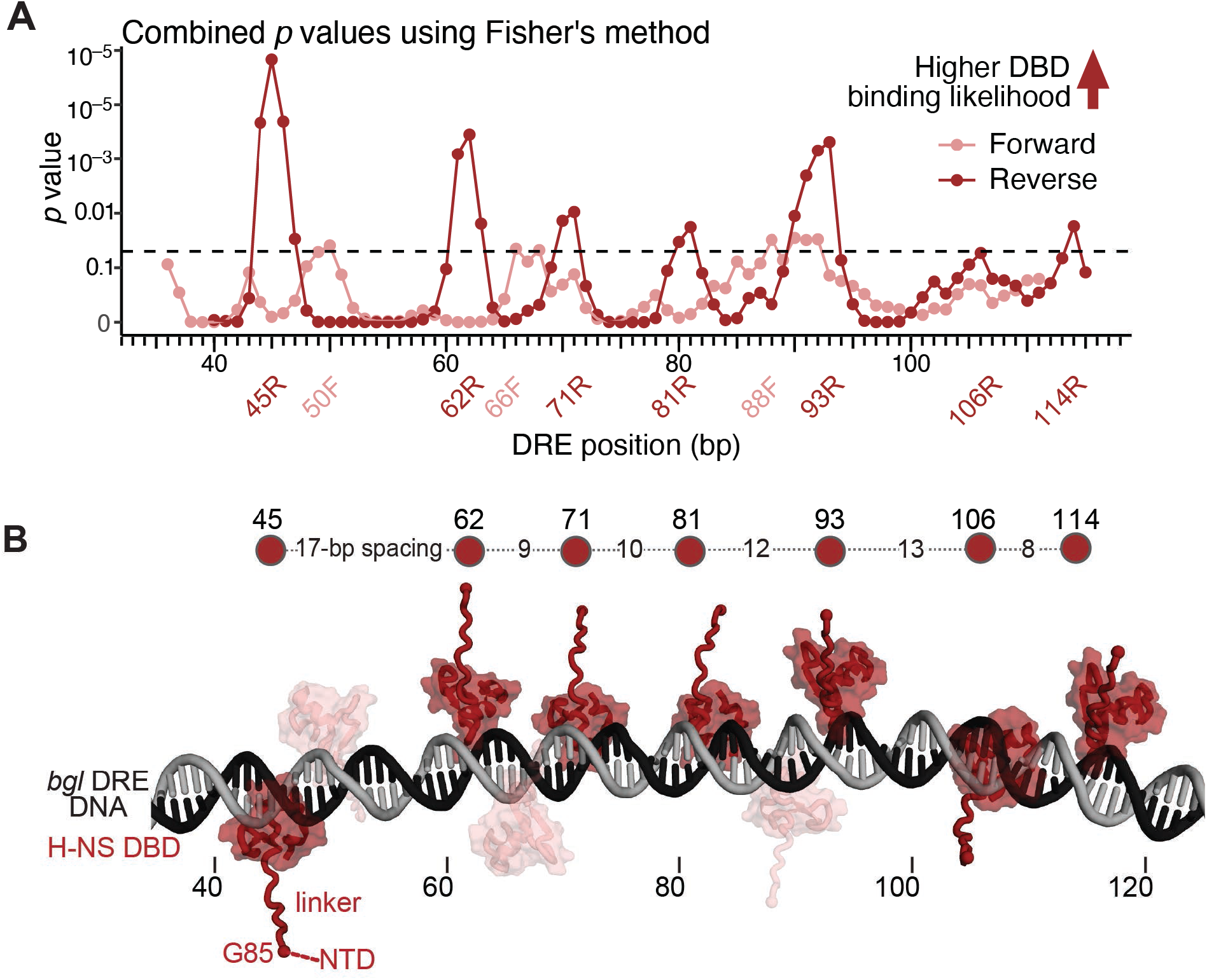
Agreement of MD simulations with TEN-map and footprinting data reveals irregularly spaced binding of H-NS DBDs to the *bgl* DRE. (A) Evidence for the localization and orientation of the H-NS DBD domain on the DRE DNA by combining evidence for congruency between the MD simulations and the TEN-map data with evidence for congruency between the MD simulations and the footprinting data (see Figure S12; dashed line, p=0.05 Fisher’s combined probability test). Evidence is shown as a combined p-value at each base pair in the DRE DNA in the forward (light red) or reverse (dark red) orientations (see Methods for details). (B) Molecular model of H-NS DBD locations in an H-NS bridged or linear filament on the *bgl* DRE. The DNA model was generated using Nucgen-plus (Bansal et al., 1995). H-NS DBDs were centered at the positions of greatest probability (panel A) in forward (light red) or reverse (dark red) orientations. The reverse orientations are most probable (see text). H-NS DBD model is from (2L93; (Gordon et al., 2011)) and the flexible linker was modeled using AlphaFold2 (Jumper et al., 2021). The spacing between DBDs in the reverse orientation is indicated on the line above the model. The H-NS NTD is not included in the model but would be attached to the DBDs via the flexible linkers.

This model of H-NS DBD interactions with the DRE DNA supports the following conclusions. First, DBD locations within the H-NS filament are irregularly spaced along the DNA duplex, with separations varying from 8-17 bp. Second, these spacings are broadly consistent with DBD contacts approximately every 10 bp along DNA possibly skipping one contact between 45 and 62, consistent with previous analyses of H-NS–DNA interactions. Third, we cannot exclude the possibility that DBDs might alternate between adjacent contact sites in the forward and reverse orientations. Finally and most importantly, the DBDs in both bridged and linear H-NS filaments do not contact DNA with rigidly fixed spacing but rather assume positions within a window of possible locations likely based on the highest affinity to local DNA sequence features (see Discussion).

### TA steps, minor groove width, and electrostatic potential best predict DBD locations

Having identified the binding locations of the H-NS DBDs on the DRE fragment, we next asked which features of the DRE DNA best predicted these locations. A prior analysis of sequences most tightly bound by the DBD of the close H-NS paralog Lsr found variable preference for AT-rich 4mer sequences (AAAA, AAAT, AATT, ATTT, TTTT, ATAA, ATAT, ATTA, AATA, TAAA, TAAT, TATA, TATT, TTAA, TTAT, or TTTA) with TA-step-containing 4mers having highest affinity when the 4mers are located in regions of high A/T content (Gordon et al., 2011). Overall AT-richness, A-tracts, or a specific high-affinity sequence (Lang motif) also have been postulated to explain H-NS–DNA binding specificity (Gordon et al., 2011; Lang et al., 2007; Navarre et al., 2006). In addition to sequence patterns, it also is possible to predict DNA structural features like minor groove width (MGW) and electrostatic potential in the minor groove (EP) (Chiu et al., 2017; Chiu et al., 2016; Zhou et al., 2013), and MGW, in particular, is suggested to favor DBD binding (Gordon et al., 2011). We first assessed the presence of these various sequence and structural features in the DRE fragment (Figure 7A). Three features, TA-step, narrow MGW, and low EP were notably enriched at sites bound by the H-NS DBD.

**Figure 7.**
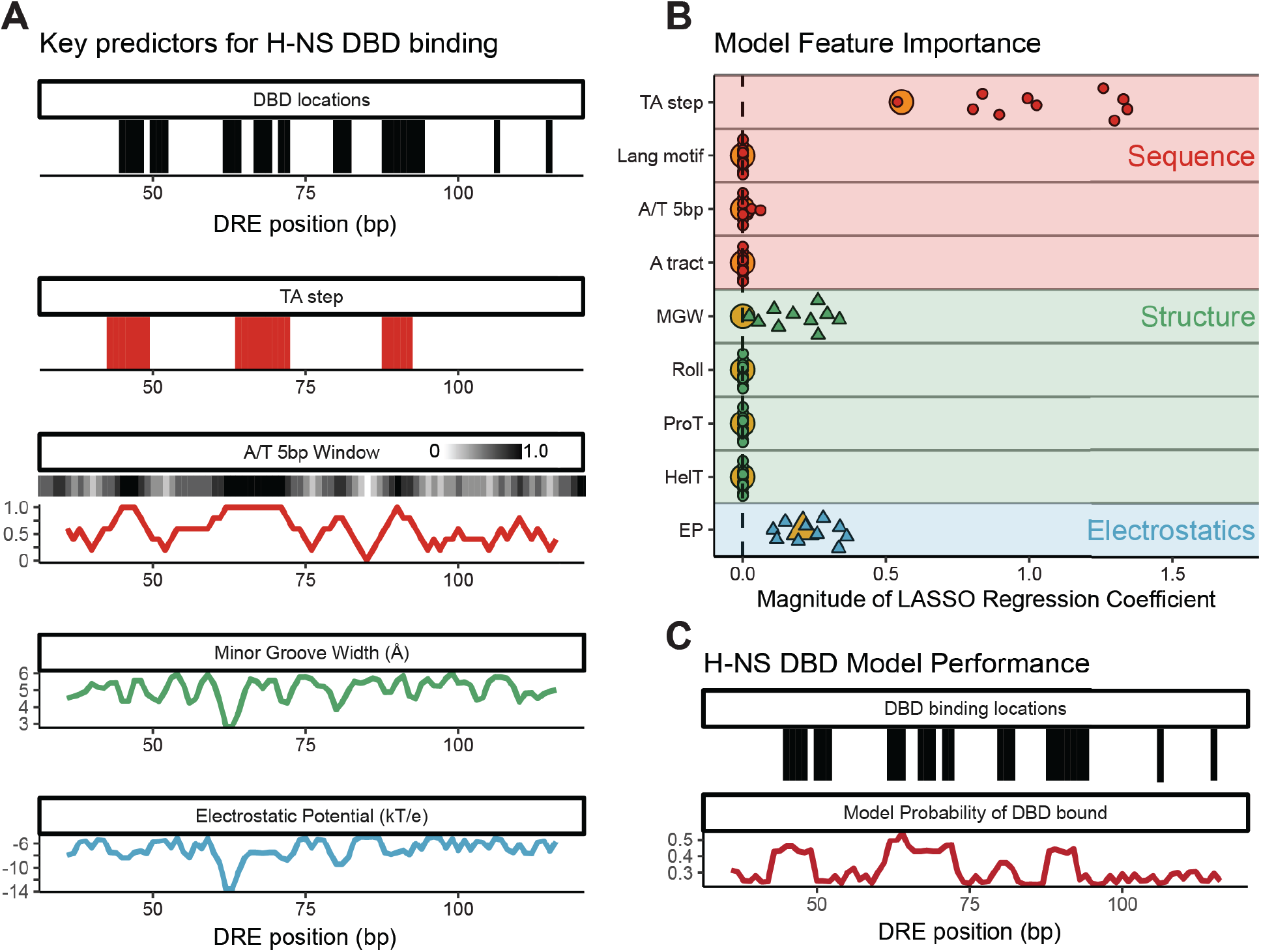
TA steps, minor groove width, and electrostatic potential best predict DBD locations. (A) Relationship between possible H-NS DBD binding locations and the sequence, structural, and electrostatic features of *bgl* DRE DNA. Base pairs with high agreement between the TEN-map and predicted patterns for footprinting and MD simulation in either DBD orientation are represented as black bars (*p* ≤ 0.05; Monte Carlo permutation test; see Methods and Figures 5C,D and S12; position 106 is included as nearly indistinguishable from *p*=0.05). TA-steps are represented as a red bar for every bp within a TA-step 4mer. A/T content averaged over a 5 bp window is shown as a white-black gradient and plot (red) above plots of minor groove width (green) and electrostatic potential (blue) calculated using DNAshapeR (Chiu et al., 2016). (B) Relative importance of sequence, structure, and electrostatic features in predicting H-NS DBD binding locations. The (+) or (-) valued regression coefficients from a LASSO logistic regression model fitted to the DBD binding locations (Friedman et al., 2010a) are represented as circles or triangles, respectively. Points correspond to coefficients from ten models fit independently by partitioning the data into ten subsets using stratified sampling and fitting each model to a unique set of nine subsets. Large orange symbols indicate the coefficients for a model fit to the entire dataset. (C) Predictive performance of sequence and structure-based regression model using TA step, minor groove width, and electrostatic potential parameters (panel B). H-NS DBD binding locations (same as in panel A) are shown in black. The probability of DBD binding based on a LASSO regression model fit to the full data set is shown in dark red.

To ask if these features were predictive of the H-NS DBD binding sites, we used a LASSO logistic regression model, which calculates the minimum number of variables required to predict an objective function, to assess the predictive contributions of sequence, structural, and electrostatic variables to the DBD binding pattern. We found that our high-confidence DRE DBD locations could be described using only TA-steps, a narrow MGW, and negative electrostatic potential (Figure 7B). As further validation that these three features can predict H-NS DBD locations, we used our regression model to predict H-NS DBD locations in the DRE fragment (Figure 7C). Encouragingly, our model was able to robustly predict the DBD binding sites that match our experimentally determined binding sites. We conclude that TA-steps, MGW, and electrostatic potential are the most useful predictors of H-NS DBD location. Since we observe the same DBD locations in both bridged and linear filaments, these features should be applicable in predicting sequence preferences for either form of the H-NS filament.

## DISCUSSION

We report new insights into the structures of bridged and linear filaments formed by the bacterial gene silencer H-NS by combining two approaches not previously applied to H-NS. First, we mapped contacts of the H-NS DBD using a tethered nuclease (TEN-map) that also causes constitutive bridging. Second, we coupled molecular and statistical analyses of TEN-map and ·OH footprinting data to predict DBD locations on a known mediator of gene silencing, the *E. coli bgl* operon DRE. These results yielded three principal insights: (*i*) HNS DBD locations do not change in the switch from linear to bridged filaments, supporting a model of DBD hemi-sequestration in linear filaments; *(ii*) the DBDs do not contact DNA with precisely fixed spacing but rather adjust both contact point and possibly orientation along adjacent minor groove positions; and (*iii*) the positions of the DBDs were best explained by the DNA sequence features AT steps, minor-groove width, and charge distribution.

These insights do not depend on complete accuracy of our DBD–DRE model (Figure 6B), which cannot be fully assessed without a high-resolution H-NS–DRE structure. Rather, our model provides a plausible way to visualize the key insights as well as the uncertainties. The constancy of irregular DBD spacing in bridged and linear filaments is our key finding. This feature of H-NS–DNA interaction remains valid even if our DBD placements are somewhat inaccurate. The irregularity and constancy of DBD interactions are compelling even though the signal-to-noise in our data is modest. Indeed, the noisy nature of H-NS–DNA interaction data, also evident in other studies, likely reflects a fundamental property of H-NS. For example, TEN-map signals for the H-NS DBDs are markedly more noisy than observed for RNAP a subunit CTDs bound to promoter DNA (Murakami et al., 1997) (compare Figure 2 to Murakami et al. Figure 5B). The H-NS DBD contacts appear to vary in strength and spacing as a function of DNA sequence for a given H-NS–DNA filament. The contacts may also be somewhat fluid. DBDs alternating in orientation, or on and off DNA, could explain why some DBD–DRE contacts appear weak or as “shadow” contacts offset by ~5 bp (Figures 6 and S12). This fluid and irregular character of DBD contacts may help H-NS vary the extent of gene silencing as a function of cellular composition, temperature, and H-NS modulators like Hha (Boudreau et al., 2018; Kotlajich et al., 2015; Qin et al., 2020; van der Valk et al., 2017; Zhao et al., 2021a; Zhao et al., 2021b).

### H-NS DBD contacts to DNA do not change in the switch from bridged to linear conformations

A key feature of H-NS-based gene regulation is the capacity of H-NS filaments to switch from binding one DNA duplex as a linear filament to binding two DNA duplexes in a bridged conformation. The extent of DNA bridging *in vivo* and whether it varies among genes or conditions remains unknown. However, the fact that H-NS can block transcript elongation only in the bridged conformation where it creates topological traps to elongating RNAP (Boudreau et al., 2018; Kotlajich et al., 2015) and the clear evidence that H-NS modulates transcription elongation *in vivo* by stimulating ρ-dependent termination (Chandraprakash and Seshasayee, 2014; Peters et al., 2012; Saxena and Gowrishankar, 2011) provide strong evidence that bridging occurs to some extent *in vivo.* A structural basis for switching of H-NS between linear and bridged filaments remains to be fully elucidated.

The Dame and Arold groups have proposed a hemi-sequestration model of H-NS DBD location that can explain linear-to-bridged switching (Shahul Hameed et al., 2018; van der Valk et al., 2017). In the hemi-sequestration model, one of the two DBDs present at each tail–tail dimerization interface in an H-NS filament binds to DNA and the second DBD is sequestered through interactions with the N-terminal dimerization domain, possibly aided by buckling of the a-helix connecting the two dimerization interfaces. Thus, the hemi-sequestered or closed conformation enables only linear filament formation whereas the open conformation, which is favored by binding of Mg^2+^ or Hha and may be constitutive in StpA (Boudreau et al., 2018; van der Valk et al., 2017), releases the sequestered set of DBDs so they can bridge to a second DNA duplex (Figure 1A; Figure 8).

**Figure 8.**
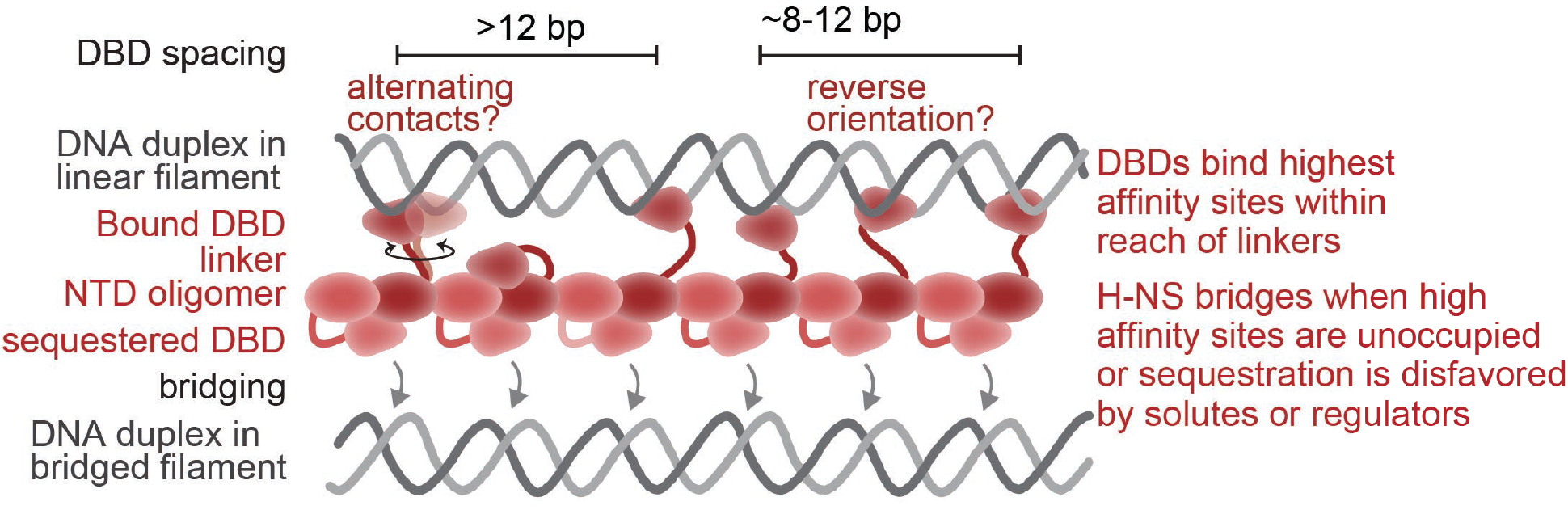
Hemi-sequestration model of H-NS DBD interactions with DNA in bridged and linear nucleoprotein filaments. In conditions favoring linear filaments, one set of DBDs is sequestered via interaction with the NTD (Shahul Hameed et al., 2018; van der Valk et al., 2017). In conditions that favor bridging, which in vivo could include altered solute composition (Qin et al., 2020), interactions of H-NS modifiers like Hha (Boudreau et al., 2018; van der Valk et al., 2017), or changes in H-NS levels (Kotlajich et al., 2015), the hemi-sequestered DBDs preferentially associate with the same set of irregularly spaced sites used in linear filaments. Variable spacing between DBDs and possible changes in DBD orientation are accommodated within the H-NS nucleoprotein filament by the flexible linkers between the DBDs and the NTD oligomer.

Our results validate the hemi-sequestration model by documenting the absence of changes in the locations of DBDs in bridged and linear filaments (Figures 3, 4, 6, and S12). This constancy of DBD contacts to DNA is evident both in the lack of effect on ·OH footprints when H-NS concentration is varied from levels known to favor bridging (≤100 H-NS/kb DNA) to concentrations far above those known to create linear filaments by saturating all available binding sites (≥200 H-NS/kb DNA; Figure 4) (Kotlajich et al., 2015). Importantly, the ·OH-footprint pattern evident in conditions favoring linear filaments predicts essentially the same DBD locations predicted for constitutively bridged filaments analyzed by TEN-map (Figure S12). This same observation of no change in footprints at different H-NS/DNA ratios or at high versus low Mg^2+^ concentration, both of which modulate linear–bridged preference, is evident in reports from several other groups (Lucht et al., 1994; Moreira et al., 2011; Ulissi et al., 2014; Will et al., 2018) (Table S1). Taken together, these results strongly favor the hemi-sequestration model of H-NS linear-to-bridged filament switching because the model readily explains use of the same DBD contacts in both bridged and linear filaments.

### DBD locations within H-NS filaments are set by DNA sequence rather than precise spacing

Our conclusion that H-NS DBD locations are unevenly spaced in H-NS nucleoprotein filaments is also consistent with previously reported ·OH and DNaseI footprints of H-NS on DNA (Lucht et al., 1994; Moreira et al., 2011; Ulissi et al., 2014; Will et al., 2018) (Table S1). These prior results suggest semi-regular spacing of ~10 bp between H-NS protected sites, but also variation in spacing over a 5–30 bp range between some sites. Our conclusion also is consistent with a single-molecule analysis of H-NS–DNA contacts (Dame et al., 2006). When bridged H-NS filaments are pulled apart with ~22 pN force, H-NS contacts are lost in steps of multiples of ~11 bp (~1 helical turn of B-form DNA). However, the ~11-bp steps are not uniform in length. Rather, they frequently deviate in apparent DBD–DNA contact positions by ± ~2 bp, consistent with our observation of DBD contacts spaced by 8–13 bp (Figure 6). Further, many ~11-bp steps give little or no detectable force signature, yielding a mean step size closer to two helical repeats. Consistent with our observation of variable strength DBD signals, some single-molecule contacts may be too weak to give a force signature or some DBDs may remain sequestered by interaction with the H-NS NTD (Shahul Hameed et al., 2018; van der Valk et al., 2017).

Consistent with this view of variable H-NS DBD contact spacing, we observed a correlation between the apparent DBD contact positions and DNA sequence features (Figure 7). Thus, DNA sequence likely dictates the lowest free energy arrangement of DBDs within the constraints imposed by their flexible tethers to the H-NS NTD oligomer. Specifically, DBDs were located on the DRE fragment at sequence and shape features that confer high-affinity for H-NS (e.g. TA steps, A-tracts leading to narrow minor groove width and negative EP), but were not observed at lower-affinity sites lacking these features (Figure 7). High affinity for TA steps also was observed in SELEX-like experiments using a DBD from an H-NS paralog (Gordon et al., 2011). Thus, the precise position and orientation DBDs on the minor groove is likely dictated by the best complement in shape and charge to the DBD within reach of the linker. Larger spacings of two or more helical repeats in contacts may reflect the absence of strong binding features near a given tail–tail junction in the H-NS NTD oligomer so that one or both DBDs at a given junction preferentially associate with the NTD.

What can this flexibility in DBD arrangement tell us about the structure of H-NS filaments? Our results define neither the specific conformation of the H-NS head–head, tail–tail oligomer nor the conformation of the bound DNA. H-NS–bound DNA is likely to deviate from a canonical B-form helix due to its AT-rich sequence and distortion by H-NS contacts. Nonetheless, our findings can guide future efforts to determine H-NS–DNA nucleoprotein filament structures and may aid understanding regulation of transcription by H-NS family proteins in several ways.

First, the DBD locations we find are inconsistent with the H-NS NTD oligomer crystal structure in which long NTD a helix separates the tail–tail junctions by ~100 Å (Arold et al., 2010). This distance would separate the junctions to which two DBD linkers connect by ≥30 bp on a B-form DNA duplex. In a bridged filament, only one of two DBDs at the junction would be available per DNA duplex. It is unlikely that H-NS can distort DNA enough to position one DBD per junction every 8-12 bp even though the 11-aa linker (G85–A95) could stretch up to ~35 Å. Other conformations of the NTD have been observed in MD simulations (Shahul Hameed et al., 2018; van der Valk et al., 2017). It seems likely that these or other alternative conformations of the NTD must explain the variable but on average ~10.5 bp DBD spacing in H-NS filaments.

Second, the irregular but sequence-dependent pattern of DBD–DNA interactions suggests that the structure of the H-NS–DNA filament itself may be irregular. Buckling of the H-NS NTD and variable extents of sequestration of DBDs via NTD interaction could both support an overall irregular structure. By allowing the DBDs to establish stronger DNA interactions with optimal sequences within range of the linker, an irregular nucleoprotein filament may allow H-NS to bind DNA more tightly than would be possible in a completely regular structure.

Finally, an irregular H-NS–DNA nucleoprotein filament structure has important implications for efforts to determine a high-resolution structure of the filament. A high-resolution structure is essential to understand H-NS regulatory function. Although cryo-EM can be used to solve structures of filaments with helically repeating structures, such approaches will not be feasible for an irregular H-NS–DNA nucleoprotein filament. Single-particle methods using H-NS bound to short DNAs such as the *bgl* DRE may be a more feasible approach.

### Limitations of the study

Our TEN-map results are limited to a single DNA fragment from the *E. coli bgl* operon for which transcriptional silencing is well characterized and our analysis of footprinting results includes only this fragment and several others from published studies. These fragments are unlikely to represent the full range of possible H-NS DBD contacts on DNAs. Further, our statistical and molecular dynamic modeling of ·OH protection and cleavage is necessarily probabilistic and does not include a full H-NS–DNA filament structure, which remains unknown. Our results thus provide a useful model that requires further testing. The challenges now are to translate this understanding to molecular resolution structures of these filament conformations and to develop approaches that can distinguish locations of linear *versus* bridged filaments in living cells.

## MATERIALS AND METHODS

### Materials

Reagents were obtained from ThermoFisher (Waltham, MA, USA) or Sigma-Aldrich (St. Louis, MO, USA), unless otherwise specified. Oligonucleotides (Table S2) were obtained from IDT (Coralville, IA); ribonucleotide triphosphates (rNTPs), from Promega (Madison, WI); [α-^32^P]GTP and [γ-^32^P]ATP, from Perkin Elmer (Waltham, MA); polyethyleneimine (PEI, avg. MW = 60,000), from Acros Organics; (S)-1-(p-Bromoacetamidobenzyl)ethylenediaminetetraacetate (BABE), from Santa Cruz Biotechnology; and DNA-modifying enzymes, from New England Biolabs (NEB; Ipswich, MA).

### Plasmid preparation

An expression plasmid for wild type H-NS (pWTHNS) was constructed by Gibson assembly using NEB HiFi reagents (Table S3). The *hns* coding sequence was amplified from *E. coli* MG1655 genomic DNA using primers #11779 and #11780 (Table S2) and the pET21d backbone was amplified from pHNS using primers #11781 and #11782 before assembly. To introduce point mutations in pWTHNS or pHisHNS (Table S3), single primer site-directed mutagenesis was performed with Q5 DNA polymerase (NEB) and primer #8964 to change the Cys21 codon (TGT) to a Ser (AGT) in pHNSC21S. To create pHNSC21SE102C, primer #8964 was used to mutate the Cys21 codon and #8963 was used to change the Gln102 codon (GAA) to Cys (TGT). Sequences of all plasmids were confirmed in key regions by Sanger sequencing.

### Protein Purification

Native StpA and Hha were purified as described previously (Boudreau et al., 2018). C-terminally His_6x_-tagged H-NS was purified as described previously (Kotlajich et al., 2015).

#### H-NS C21S purification

H-NS C21S was purified as described for non-tagged H-NS in (Boudreau et al., 2018) with the following modifications. Briefly, BL21 λDE3 cells transformed with pHNSC21S and pHiC were grown in 2 L of LB supplemented with ampicillin (100 μg/mL) + gentamycin (10 μg/mL) to OD_600_ ~0.6 before inducing expression of H-NS C21S with 0.5 mM isopropyl β-D-1-thiogalactopyranoside (IPTG) for 2 h at 37 °C. Pelleted cells were lysed by sonication. H-NS C21S was precipitated with addition of 0.6% (w/v) PEI and eluted with PEI elution buffer containing 0.8 M NaCl followed by ammonium sulfate precipitation. Precipitated H-NS was dissolved in nickel binding buffer and applied to a HisTrap Ni^2+^ column (GE Healthcare). Fractions containing H-NS were mixed with TEV protease to remove the N-terminus His_6x_-tag. H-NS C21S was reapplied to the nickel column. Non-tagged protein was collected in the flow through and dialyzed against Heparin Buffer (20 mM Tris-HCl pH 7.5, 130 mM NaCl, 100 μM EDTA, 5% glycerol, 1 mM dithiothreitol (DTT)). H-NS C21S was then applied to a HiTrap Hep column (GE Healthcare) and eluted with Heparin Buffer containing 1 M NaCl. Fractions containing H-NS C21S were pooled and dialyzed against Binding Buffer (20 mM Tris-HCl pH 7.2, 10% glycerol, 1 mM DTT) supplemented with 130 mM NaCl. To concentrate H-NS, the protein was applied to a Resource Q column (GE Healthcare). The column was washed with a 130–200 mM NaCl gradient in Binding Buffer over 7 column volumes. H-NS C21S was eluted in 3 mL Binding Buffer with 500 mM NaCl. Fractions containing H-NS were dialyzed against H-NS Storage Buffer (20 mM Tris-HCl pH 7.5, 0.3 M potassium acetate, 10% glycerol) before storing at −80 °C. Protein purity (~90%) and yield (~5 mg) were determined by SDS-PAGE and Qubit, respectively.

#### Native WT and C21S E102C H-NS purification

BL21 λDE3 cells were transformed with pWTHNS or pHNSC21SE102C before deleting *hns* by P1 transduction using a lysate from the Δ*hns::kan* Keio strain (Coli Genetic Stock Center #9111; (Baba et al., 2006). After confirming the *hns* deletion by PCR, 2 L of cells were grown aerobically in Luria Broth–0.5% NaCl supplemented with 100 μg ampicillin/mL at 37 °C to apparent OD_600_ 0.5 before inducing H-NS expression by addition of IPTG to 1 mM and carbenicillin to 250 μg/mL followed by shaking at 16 h at 25 °C. Cells were pelleted and resuspended in 40 mL of lysis buffer (20 mM Tris-HCl, pH 7.2, 130 mM NaCl, 10% glycerol, 1 mM DTT, 1 mM β-mercaptoethanol supplemented with phenylmethylsulfonyl fluoride (PMSF) and protease inhibitor cocktail (PIC; 31.2 mg benzamide/mL, 0.5 mg chymostatin/mL, 0.5 m leupeptin/mL, 0.1 mg pepstatin/mL, 1 mg aprotonin/mL, 1 mg antipain/mL). Cells were lysed by sonication and the lysate was cleared by centrifugation at 27,000 × *g* for 45 min at 4 °C. H-NS containing cell lysate was directly applied to a HiTrap Heparin HP column (GE Healthcare) equilibrated with Binding Buffer with 130 mM NaCl. H-NS was eluted at ~600 mM NaCl using a gradient elution from 130 mM–1 M NaCl over 20 column volumes. Fractions containing H-NS were pooled and dialyzed against binding buffer containing 130 mM NaCl. H-NS was applied to a 1-mL Resource Q column equilibrated with Binding Buffer+130 mM NaCl. Remaining impurities were eluted over a gradient of 130–200 mM NaCl in 10 mL of Binding Buffer. H-NS was eluted with 3 mL of Binding Buffer+500 mM NaCl. Fractions containing H-NS were dialyzed against H-NS Storage Buffer supplemented with 5 mM TCEP. Purity (~90%) and amount of protein (~4 mg) was determined by SDS-PAGE and Qubit analysis, respectively. For tethered cleavage or ·OH cleavage assays, wild-type H-NS was exchanged into a 3x stock of Filament Buffer (3x concentrations: 120 mM HEPES-KOH, pH 8.0, 300 mM potassium glutamate, and 15% glycerol).

### FeBABE Conjugation

H-NS C21S and H-NS C21S E102C were exchanged into Conjugation Buffer (10 mM MOPS, pH 8.0, 0.5 mM EDTA, 200 mM NaCl, 5% glycerol) using PD SpinTrap G-25 columns (GE Healthcare #28-9180-04) pre-equilibrated with conjugation buffer. FeBABE was charged by incubating (S)-1-(p-Bromoacetamidobenzyl)ethylenediaminetetraacetate (BABE; Santa Cruz Biotechnology) with FeCl_3_•6H_2_O as previously described (Greiner et al., 1997). H-NS variants (50 μM) were incubated with 10-fold molar excess FeBABE for 2 h at room temperature to conjugate FeBABE to H-NS. After conjugation, 7.6 μmol of conjugated H-NS was combined with 20-fold-molar excess mPEG24 (ThermoFisher # 22713) for an additional 2 h at room temperature. Samples were visualized on 10% Bis-tris gel (ThermoFisher) to determine the percent conjugated (~50% for H-NS C21S E102C). From the remaining conjugation solution, excess FeBABE was removed by applying conjugated H-NS to a clean PD SpinTrap G-25 column equilibrated with a 3X stock of Filament Buffer. Conjugation was also confirmed by the appearance of a second, slower migrating band on a 20% polyacryamide PhastGel (GE Healthcare). Fe^2+^·EDTA–H-NS was stored in aliquots at −80°C and remained stable under these storage conditions for up to 1 year. The DNA-binding activity of the conjugated protein was confirmed by electrophoretic mobility shift assay and by *in vitro* transcription assays described below.

### In vitro transcription assay

In vitro transcription of the *bgl* nucleoprotein filament was preformed essentially as described previously (Boudreau et al., 2018; Kotlajich et al., 2015) using 1.5 kb DNA fragment generated by PCR from pMK110 (Haft et al., 2014; Kotlajich et al., 2015) with PCR primers 645 and 3071 (Table S2). In brief, 150 nM *E. coli* RNAP (Gribskov and Burgess, 1983; Nayak et al., 2013) reconstituted into holoenzyme with σ^70^ was incubated with a 100 nM DNA containing the λ_Pr_ promoter fused to *bglF–bglG* DNA to form halted elongation complexes (A26 ECs) in 40 mM HEPES-KOH, pH 8.0, 100 mM potassium glutamate, 8 mM magnesium aspartate, 0.022% NP-40, 100 μg acetylated bovine serum albumin/mL, 10% glycerol, 1 mM DTT, 150 μM ApU, 10 μM each ATP and UTP, 2.5 μM [a-^32^P]GTP for 15 min at 37 °C. The halted A26 ECs were then diluted to 10 nM in the same solution plus 100 μg rifampicin/mL and 0.1 U RNasin/μL and incubated with wild-type or modified H-NS at the concentrations indicated in the figures for 20 min at 20 °C to form nucleoprotein filaments. Transcription through the filaments at 20 °C was then restarted by adjusting to 4 NTPs to 30 μM each final. Samples were removed at the times indicated, extracted with phenol, ethanol precipitated, and electrophoresed through a 6% polyacryalmide gel (acrylamide:bisacrylamide 19:1) in 8 M urea, 44 mM Tris borate, pH 8.3, 1.25 mM EDTA, imaged using a GE Life Sciences Typhoon phosphorimager, and converted to pseudo-densitometry plots as described previously.

### Electrophoretic mobility shift assay

EMSA was performed as described previously (Boudreau et al., 2018; Kotlajich et al., 2015). Briefly, 1.5 kb *bgl* DNA fragments, which were the same sequence used in in vitro transcription assay, were radiolabeled during PCR. DNA was then incubated with various H-NS proteins as indicated at 20 °C to form filaments. Filaments were run on a using 3% polyacrylamide gels at 4 °C and imaged using a GE Life Sciences Typhoon phosphorimager.

### Tethered-nuclease mapping (TEN-map) Assay

The 146-bp DRE DNA for the TEN-map assay was generated by PCR from a plasmid pMK110 using primers #7486 and #12638 (Tables S2 and S3). To enable visualization by phosphorimaging, one primer was 5’-end-labeled with [γ-^32^P]ATP prior to the PCR reaction. The PCR product was separated on 5% non-denaturing PAG, eluted from the gel overnight at 4 °C in 20 mM Tris-HCl pH 7.5, 1 mM EDTA, and 200 mM NaCl, and concentrated using QIAquick gel extraction kit. Increasing concentrations of Fe^2+^·EDTA–H-NS (from 50 to 5000 nM) were incubated with ~18 nM DRE DNA for 20 min at 20 °C in 7.5 μL of 1X Filament Buffer supplemented with 8 mM magnesium aspartate. Hydroxyl radical cleavage was initiated by addition of 1 μL 2 mM H_2_O_2_ and 1 μL 2 mM ascorbic acid. Cleavage proceeded for 2 min at 20 °C after which 8 μL of the cleavage reaction was added to 100 μL TE-saturated phenol, pH 7.9 and 92 μL cleavage stop solution (20 mM EDTA, 20 μg glycogen). The cleavage products were recovered by phenol extraction and ethanol precipitation, and then resuspended in 4 μL formamide stop dye (95% formamide, 20 mM EDTA, 0.02% xylene cyanol, 0.02% bromophenol blue). The samples were electrophoresed through a 7% urea polyacrylamide gel (40.4 x 45.7 cm) in 0.5X TBE and alongside a Maxam-Gilbert A+G ladder for alignment (Maxam and Gilbert, 1977) and MspI-digested, ^32^P-end-labeled pBR322 for size reference. The cleaved DNA was quantified using the densitometric, semi-automated footprinting analysis (SAFA) software (Das et al., 2005). The resulting peak areas were first normalized to total signal in each sample as determined by ImageQuant (version 5.2; GE Lifesciences) analysis of an unsaturated phosphorimage scan of gel and then adjusted to values above and below the average peak area in each sample.

For the offset DNA experiment (Figure S5A-B), the TEN-map assay was performed as described above except using a 151-bp DRE DNA amplified from pMK110 using 5’ ^32^P-end-labeled primer #14374 and unlabeled primer #12386. For the TEN-map assay on the 549-bp DRE fragment (Figure S5C-D), DNA was amplified from pMK110 using 5’ ^32^P-end-labeled primer #7486 and unlabeled primer #7491. A mixture of Fe^2+^·EDTA–H-NS and wild-type H-NS at 4:6 ratio and concentration indicated in the figure legend was added to ~18 nM DNA in a 20 μL reaction in Filament Buffer supplemented with 0.02% NP-40 and 100 μg acetylated BSA/mL (supplemented Filament Buffer) to compare to the bridged filament cleavage described below. Cleavage was initiated by addition of 1 μL 50 mM ascorbic acid and 4 μL 12.5 mM H_2_O_2_. Cleavage continued for 2 min at RT before stopping the reaction with 25 μL 22.4 mM thiourea and 20 μL TE. The cleavage products were recovered by phenol extraction and ethanol precipitation, and then resuspended in 4 μL formamide stop dye. The cleavage products were visualized and quantified as described above for the 146-bp DRE DNA. To test the effect of Mg^2+^ on the H-NS DBD location and orientation (Figure 3B and S8), the TEN-map assay was performed on the 549 bp DNA fragment as described except the amount of magnesium aspartate was varied as specified in the figure legend.

### Hydroxyl radical footprinting

H-NS filaments were formed on the same 146-bp DRE DNA template used in the TEN-map assay. Concentrations of unmodified H-NS from 50 nM to 4 μM (17–1360 H-NS/kb) were incubated with ~20 nM 5’ ^32^P-end-labeled DRE DNA in 50 μL reactions in Filament Buffer without glycerol. Filaments were allowed to form at 20 °C for 20 min prior to ·OH cleavage. Cleavage was initiated by addition of 0.6 μL 100 mM ascorbic acid, 4 μL 0.15% H_2_O_2_, and 6 μL fresh 50 mM Fe^2+^-EDTA (50 mM (NH4)2Fe(SO4)2, 100 mM EDTA) and incubated for 2 min at 20 °C. The no-cleavage control was treated identically except 10.6 μL water was added in place of cleavage reagents. ·OH was then quenched by addition of 80 μL 20 mM thiourea and 60 μL of 0.3 M NaCl, 1 mM EDTA. Cleavage products were recovered by phenol extraction and ethanol precipitation, and then resuspended in 4 μL formamide stop dye. Samples were visualized as described for TEN-map assay and quantified using SAFA (Das et al., 2005).

### Bridged H-NS cleavage assay

Bridged H-NS filaments were formed using the previously published bridging pull-down assay (van der Valk et al., 2018; van der Valk et al., 2017) with the following modifications. The 5’ ^32^P-end-labeled, 549-bp DRE DNA (see above) was used as the prey DNA. To form bridged filaments, 5 nM biotinylated 685-bp DNA fragment (68% AT; Table S2) coupled to 3 μL M280 streptavidin dynabeads (ThermoFisher) was incubated with 5 nM prey DNA and H-NS in supplemented Filament Buffer at 5–8 μM 6:4 WT H-NS: Fe^2+^·EDTA–H-NS as indicated in the legend to Figure 3. High levels of H-NS (>5 μM) are required to form bridged filaments in the bridging/pull-down assay, possibly because DNA sequestration on the positively charged streptavidin beads inhibits H-NS binding (Figure 3A; (van der Valk et al., 2017). Bridged filaments were formed for 20 min at 20 °C with shaking at 1000 rpm in an Eppendorf Thermomixer and then separated from unbound protein and DNA using a magnetic rack. Bead-immobilized bridged filaments were then resuspended in 20 μL supplemented Filament Buffer followed by addition of 1 μL 50 mM ascorbic acid and 4 μL 12.5 mM H_2_O_2_. Cleavage was proceeded for 2 min at RT before addition of 25 μL 22.4 mM thiourea and 20 μL TE. DNA recovered and resuspended in formamide stop dye as described above. The amount of ^32^P retained on the beads was measured by counting Cerenkov radiation and compared to amounts retained using only unmodified H-NS and prey DNA. Bridging efficiency was calculated as a fraction of total prey DNA ^32^P that was retained on the beads. Apparent bridging efficiency was likely decreased by some DNA lost during sample processing. Cleavage products were then visualized and quantified by gel electrophoresis and phosphorimaging as described above for the TEN-map assay.

### MD simulation of FeBABE-H-NS bound to DNA

Molecular dynamics simulations were run using the program OpenMM (Eastman et al., 2017) with the Charmm36 force field (Best et al., 2012; Hart et al., 2012). Initial setup and post-processing analyses were conducted using the program CHARMM (Brooks et al., 2009). We obtained an initial starting model of H-NS bound to a 12-bp strand of duplex DNA from Vreede and coworkers (Gordon et al., 2011). We “mutated” the side-chain of Cys102 in that model to an FeBABE moiety and generated force field parameters for the organic component of FeBABE by analogy to existing functional groups in the Charmm36 force field. We modeled the Fe^2+^ using a cationic dummy-atom model (Liao et al., 2017). We dissolved the H-NS/FeBABE/DNA system in a cubic box of water with 70 Å edges and added Na^+^ and Cl^-^ ions to reach an ionic strength corresponding to 150 mM with neutral overall charge. The complete system contained 31,518 atoms. Following a short geometry optimization, the system was heated from 48 K to 298 K during 1 ns and equilibrated at 298 K and 1 atm for an additional 4 ns prior to production simulations. The simulations employed the NPT ensemble using the Andersen thermostat and MC barostat. Non-bonded interactions were cutoff above a distance of 12 Å with a switching function from 10 Å to 12 Å and the integration time step was 1 fs. We placed a harmonic restraint with force constant of 1 kcal/mol•A^2^ on DNA backbone atoms to prevent unraveling at the ends of the duplex, which we observed in initial tests. Under these conditions, we simulated the system for 500 ns, taking snapshots every 10 ps.

In each frame, we calculated the distance from the Fe^2+^ to every C5’, because FeBABE-induced cleavage is thought to occur by ·OH radical reacting with C5’-H (Balasubramanian et al., 1998). We modeled the concentration of ·OH radicals at C5’ based on a diffusion model where the concentration of radicals around the Fe^2+^ decays according to a Gaussian distribution (Cheal et al., 2009). To account for cleavage sites that are shielded from solvent by the bound protein, the average concentration of ·OH at each site was scaled linearly by the average solvent accessible surface area of the corresponding C5’ methylene group (Figure 5B). The scaled concentration at each site yields the cleavage propensities displayed in Figure 5A. The simulations used a relatively short 12-bp duplex in order to achieve reasonable computational efficiency and to run simulations long enough to adequately sample the conformational flexibility of the FeBABE. To model cleavage propensities for bp at greater distances from the H-NS binding site, we assumed that long-range interactions with DNA do not influence the overall behavior of FeBABE and thus estimated those cleavage propensities by adding additional DNA to the saved trajectories after the simulation was complete.

### Statistical modeling of DBD locations in H-NS filaments

The ·OH footprinting data for each H-NS concentration tested (0–4 μM titration), each strand, and each independent replicate was normalized by dividing the signal in each lane by the average signal within a lane. The resulting ratio was log_2_ transformed to center the data at zero. The log_2_-normalized deviance from the average signal from each strand was separately fit to a Bayesian multi-level model of the following form:

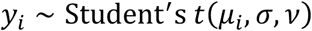

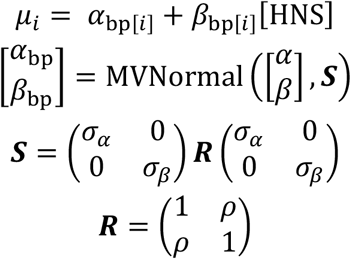

With the following weakly regularizing priors:

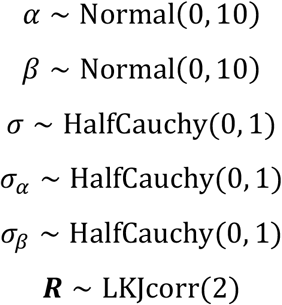

The models were fit using the R package brms (version 2.10.0) (Bürkner, 2017) and default priors were used for any parameter for which a prior is not specified above.

Similarly, the tethered-cleavage data for each concentration (0–3 μM titration) was normalized by first dividing each lane by a constant factor resulting from quantification of a short scan of the entire lane. The data was then centered by subtracting the average signal within each lane. The centered and normalized signal for each strand separately was then fit to a Bayesian multilevel, of the form described above, with the following priors:

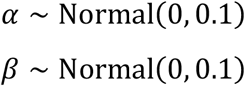

Default priors were used for every parameter not specified above. Notably, the 4 μM and 5 μM concentrations were not included in the tethered cleavage fits due to their substantial deviance from linearity. Point estimates for each slope at each bp in the models above were estimated from the average of the posterior distribution for that slope. These estimates can be interpreted as the H-NS concentration-dependent change in signal at each bp with data-driven shrinkage towards the grand mean of slopes for every bp.

To match the MD simulations with the *in vitro* data, the slope estimates for each strand in each data type were Robust-Z normalized using the following formula:

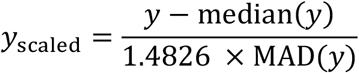

where MAD is the median absolute deviation and 1.4826 is a scaling factor to convert the MAD to an estimate for the standard deviation (Hampel, 1974; Rousseeuw and Croux, 1993). Similarly, the expected 15-bp cleavage pattern of a single H-NS DBD (Figure 5A) was also scaled to a standard Normal using the formula above. Using a sliding window, the Manhattan distance between the scaled, tethered-cleavage estimates and the scaled, 15-bp expected cleavage pattern was calculated for each base centered on the sixth bp of the cleavage pattern (triangle in Figure 5). Due to the asymmetry of the expected cleavage pattern, we performed this calculation in both directions along the DNA corresponding to two different possible orientations of the H-NS DNA binding domain. Since this score is a distance, lower values indicate a closer match between the expected cleavage pattern and the tethered cleavage data.

To assess the statistical significance of the MD match scores, we performed a one-sided Monte Carlo permutation test by considering the MD match score at each base pair in 1000 different random shuffles of the tethered-cleavage data for each direction of the H-NS DBD. We generated a *p*-value by determining the fraction of shuffled datasets that had a match score greater than or equal to the match score with unshuffled data. We considered MD match scores below the 5^th^ percentile of the random MD match scores (*p*-value < 0.05) to signify a statistically significant match between the expected cleavage pattern and the tethered cleavage data.

To determine possible localizations of the H-NS DBD in the ·OH footprinting data, we performed the same procedure described above using the solvent accessibility of the 5’ carbon of the DNA duplex as the expected pattern of protection imparted by the H-NS DBD (Figure 5B) and matched that pattern to the ·OH footprinting data (Figure 5F,H).

To combine evidence for the DBD locations between the footprinting and TEN-map datasets (Figure 6) we used Fisher’s method (Heard and Rubin-Delanchy, 2018) to generate combined *p*-values for each base pair and DBD orientation.

### Predicting H-NS DBD binding locations

To predict H-NS DBD binding a dataset consisting of sequence, structural, and electrostatic parameters was generated. **Sequence** – For each bp, the following sequence features were generated: the A/T content of a 5bp window centered on the bp (A/T content-5p), overlap with a 4mer matching the criteria for an AT tract, overlap with a 4mer matching the criteria for a TA step, and overlap with a motif match to the Lang et al. H-NS PWM (Lang et al., 2007). Motif matches were generated by running FIMO (version 5.05) (Grant et al., 2011) over the *P-bgl* fragment using a third-order Markov background model trained on the *E. coli* genome, searching on both strands, and using a *p*-value cutoff of 0.001 with no FDR correction. **Structure and electrostatics** - DNA structure and electrostatic features were generated using the DNAshapeR package (Chiu et al., 2016) using default parameters on the *P-bgl* fragment.

To predict the locations of H-NS DBD binding at the base-pair level, a LASSO logistic regression model was trained on the H-NS TEN-map and footprinting data using glmnet version 2.0-16 (Friedman et al., 2010b). For these models, we assigned as bound by H-NS base pairs for which the *p*-value for a match to the predicted patterns (Figures 5C,D and S12) was less than 0.05 for either the TEN-map or the footprinting data in either orientation (position 106 was excluded from the LASSO model). All continuous features were centered by their average and scaled by their standard deviation before training. The LASSO model reveals the minimal set of DNA features needed for the prediction of a given dataset by only including non-zero coefficients for a feature if that feature provides independent information from other non-zero coefficients already present in the model. Since all input data was centered and scaled before training the model, the relative magnitude of each coefficient can be interpreted as the importance for that feature in predicting the outcome, with positive values indicating an increase in the probability of a region being bound by a protein when the associated feature is a higher value and negative values indicating the opposite. Models were trained using 10-fold cross-validation with stratified sampling and the penalty parameter, λ, set as one standard error away from the minimum binomial deviance. λ was determined individually for each subsampled dataset using 10-fold cross-validation internally to test for sensitivity to the exact data used to train the model. Predictive performance was assessed using a model trained on the full data set with 10-fold cross-validation used to determine the λ parameter.

## ACKNOWLEDGEMENTS

We thank Jocelyne Vreede (University of Amsterdam) for sharing coordinates for an H-NS DBD–DNA structure and for modeling FeBABE into the structure. We thank members of the Landick lab for many helpful discussions and comments on this manuscript. This work was supported by NIH grant GM38660 to R.L. B.A.S. was supported by an NSF Graduate Research Fellowship (DGE-1256259). C.M.H. was supported by an NSF Graduate Research Fellowship (DGE-1747503) and an NIH Chemical Biology Interface Training grant (T32 GM008505). Computational resources for MD simulations were provided by the Extreme Science and Engineering Discovery Environment (XSEDE; allocation TG-MCB180084 to D.R.), which is supported by National Science Foundation grant number ACI-1548562.

## Select Abbreviations

DBD: DNA binding domain
DBP: DNA binding protein
FeBABE: Fe(III) (S)-1-(p-Bromoacetamidobenzyl)ethylenediaminetetraacetate
Fe^2+^·EDTA–H-NS: H-NS derivatized with an Fe^2+^·EDTA nuclease
MD: molecular dynamics
NTD: N-terminal domain
·OH: hydroxyl radical
RNAP: RNA polymerase

