## Supplemental material for "Bacterial H-NS contacts DNA at the same irregularly spaced sites in both bridged and hemi-sequestered linear filaments"

**Table S1. Reported H-NS footprinting patterns for diverse DNA fragments**
Reported H-NS footprinting patterns from studies in which H-NS concentrations were varied and experiments conditions could be ascertained. Description of experimental conditions used in previously described H-NS footprinting experiments. Features of H-NS protected regions such as the length of the protected regions, spacing between protected regions, and any changes as a function of H-NS/kb ratio are described.

| | Publication | (Lucht et al., 1994) | (Moreira et al., 2011) | (Ulissi et al., 2014) | (Will et al., 2018) |
|---|---|---|---|---|---|
| Experimental conditions | Method | DNaseI footprinting | DNaseI footprinting | DNaseI and ·OH footprinting | DNaseI Differential DNA Footprint Analysis |
| | DNA sequence | *E. coli proU* promoter | *E. coli bolA* promoter | *Shigella virF* promoter | *Salmonella pagC* promoter |
| | DNA length | (i) 256 bp<br>(ii) 445 bp<br>(iii) 262 bp | 310 bp | 450 bp | 8.1 kb linearized plasmid |
| | H-NS/kb DNA (range) | 111–9687 (i,iii)<br>67-5573 (ii) | 483–4193 | 12.5–125 (H-NS); 7,000–113,000 (DBD only) | 2.5–125 |
| | $Mg^{2+}$ concentration | 15 mM $Mg^{2+}$ | 10 mM $Mg^{2+}$ | 1.5 mM $Mg^{2+}$ | 0–50 mM $Mg^{2+}$ |
| | *Filament* type *(if known)* | Unknown | Unknown | Unknown | Bridged (low H-NS/kb)<br>Linear (high H-NS/kb) |
| Features of protected regions | Average length | 20 bp | 19.2 bp | 8.3 bp (FL); 7 bp (CTD only) | <5 bp |
| | Spacing | Median is ~ 7 bp ; varies from 2 - 17 bp | Median is 17 bp; varies from 15-32 | Some ~10 bp spacing; varies from 3-20 bp | Median 13 bp; varies from 5-21 bp |
| | Changes as a function of H-NS/kb? | No change in location, but more robust protection at >1000 H-NS/kb | No change in location, but more robust protection at >3,000 H-NS/kb | No change in location | No change in location, but more robust protection at >25 H-NS/kb |

Table S2. Oligonucleotides used in this study.

| Primer Number | Description | Sequence |
|---|---|---|
| 8964 | C21S site-directed mutatgenesis | 5′CGTGCGCAGGCAAGAGAAagtACACTTGAAACGCTGGAAGAAATGC |
| 8963 | E102C site-directed mutatgenesis | 5′CCGGCAAAATATAGCTACGTTGACtgtAACGGCGAAACTAAAACCTGG |
| 11781 | pET21d_rev | 5′–TGAGATCCGGCTGCTAAC |
| 11782 | pET21d_fwd | 5′GGTATATCTCCTTCTTAAAGTTAAACAAAATTATTTCTAGAGGGGAATTGTTATCCGC |
| 11779 | wtHNS_fwd | 5′–ttgttagcagccggatctcaTTATTGCTTGATCAGGAAATC |
| 11780 | wtHNS_rev | 5′–CtttaagaaggagatataccATGAGCGAAGCACTTAAAATTC |
| 12638 | DRE NT strand | 5′–CTGGTCAGTGCCCAAATGAG |
| 7486 | DRE Temp strand | 5′–AGATGTGTAACCAGTCGCTGA |
| 14374 | DRE Temp offset | 5′-ACTTCAGATGTGTAACCAGTCG |
| 13444 | Bait_fwd_biotin | 5′/5Biosg/ATACATATGCAACTTGAACGGCGTAAAAGAGGAACAATGG |
| 13445 | Bait_rev | 5′-GGTGGATCCTTTTCATCCCTTTAGTTCTTCCAG |
| 1343 | BaitDNA_geneblock (double stranded DNA) | 5′ATACATATGCAACTTGAACGGCGTAAAAGAGGAACAATGGAAATAATGTTTGATATATTGAGGAATTGTGAGCCAAAATGCGGAATAACGAGAGTAATATACGGTGCTGGTATAAATTATGTAGTAGCTCAAAAATATTTGGATCAATTGGTAAAAGTAGGTGCTTTAAACATTAAAACTGAAAATGATAGAAAGATTTATGAGATAACTGAAAAGGGTAAGTTACTAAGGACTCATATCGAAGAATTCATAAAGATAAGAGAGAATTTGTATTCGGCTAAAGAGAAAGTTAGTGAACTTTTAAGAACAGACAGTGAGTAAACAAGGTGTAGCTAGAGTTTTCTCTTTATTCTTTCTAGTCGAATTTTCTCTTCAAATGTTTTCTCCAATTGATTTAGTCTATTCGATATATGGACTCTTAGTGGCAAATATATAATGTATAAGTTCTGGAGTAGTATGGTAATAACTATTTTATTTTCGATAGCTTGAATGTTTATTTTCCCAGAGACATTAGGTGGTCTTTCAAACTCAATAATGTGGGTAATATTATAAGATCCTACATATTCTCTACCTTGGAAAACAACCGGTAATTTCCCAATATCGCCTTCACATCTATAGAGTCTACTATCAGAGTTTAGGAGAAATTTTGTAACTGGAAGAACTAAAGGGATGAAAAGGATCCACC |
| 7491 | DRE long rev | 5′–GTACGAATTCGGAATTGGCTTTCAAAAACG |
| 645 | *bgl*_fwd | 5′-CAGTTCCCTACTCTCGCATG |
| 3071 | *bgl*_rev | 5′-CGTTAAATCTATCACCGCAAGGG |

Table S3. Plasmids used in this study.

| Plasmid Name | Plasmid Number | Description | Reference |
|---|---|---|---|
| pHNS | 4548 | *E. coli* H-NS with N-terminus 6xHis tag and TEV cleavage site downstream of IPTG inducible promoter | (Boudreau et al., 2018) |
| pHNSC21S | 5049 | *E. coli* H-NS C21S point mutation with N-terminus 6xHis tag and TEV cleavage site downstream of IPTG inducible promoter in pET21d (+) backbone | This work |
| pWTHNS | 5052 | *E. coli* H-NS coding sequence cloned without the His-tag downstream of IPTG inducible promoter in pET21d (+) backbone | This work |
| pHNSC21SE102C | 5055 | Modified version of pWTHNS with C21S and E102C point mutations generated by site-directed mutagenesis with primers #8964 and #8963, respectively | This work |
| pHiC | | Single-copy BAC plasmid carrying an engineered $lacI^{q1}$ repressor allele that expresses approximately 170-fold more Lac repressor than the wild-type *lacI* gene. Isolated from Lucigen strain HI-Control 10G. | Lucigen |
| pMK110 | 5605; Addgene ID #99535 | Antisense *bgl* operon downstream of lambda $P_R$ promoter and 26 bp C-less cassette used for generation of *in vitro* transcription templates or DNA substrate for H-NS filament formation in TEN-map and hydroxyl radical footprinting experiments | (Haft et al., 2014; Kotlajich et al., 2015) |

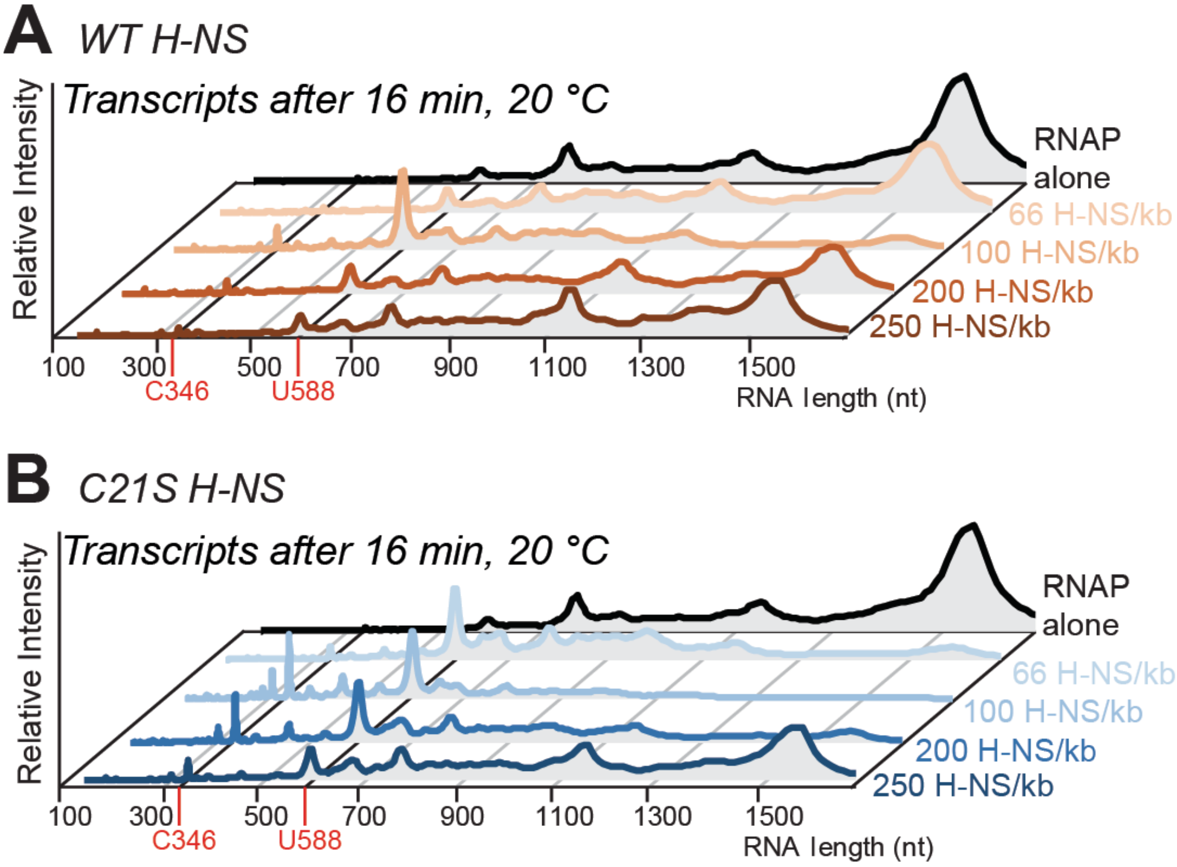


**Figure S1. Effect of WT H-NS and C21S H-NS on an elongating RNAP.**
Pseudo-densitometry traces of transcripts formed on a 1.5 kb *bgl* DNA by RNAP 16 min after transcription through various H-NS filaments is restarted with addition of NTPs (30 μM).
(A) WT H-NS filaments.
(B) C21S H-NS filaments.
H-NS–DNA filaments switch from bridged to linear conformations at ≥200 H-NS/kb. DNA (see Figure S2).

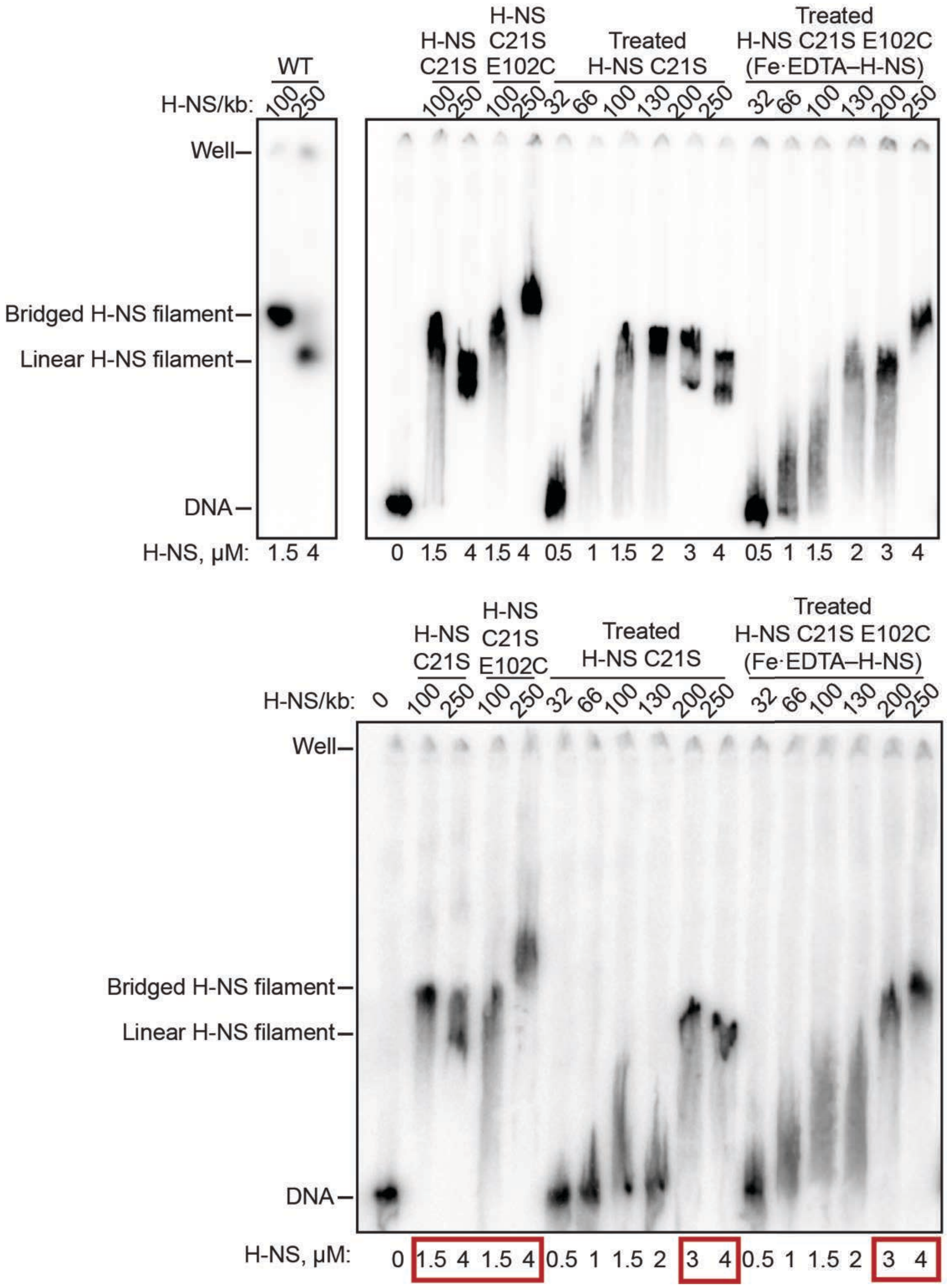


**Figure S2. $Fe^{2+}$·EDTA–H-NS forms filaments on *bgl* DRE DNA.**

EMSA of H-NS filaments formed at 10 nM DNA on the 1.5 kb *bgl* DNA fragment used for *in vitro* transcription assays (see Figs. 11E,1F and S1) (Boudreau et al., 2018; Kotlajich et al., 2015). Filaments were formed with WT H-NS, H-NS C21S, or H-NS C21S E102C with or without FeBABE treatment. H-NS C21S was treated with FeBABE to test for loss of activity during conjugation process, but H-NS C21S was not derivatized by FeBABE because it lacks reactive thiols. $Fe^{2+}$·EDTA–H-NS contains FeBABE conjugated to ~50% of the E102C thiols in H-NS C21S E102C (see Methods). Top and bottom panels depict filament formation by two independent preparations of $Fe^{2+}$·EDTA–H-NS. The order of lanes in the top panel was edited for clarity. Red boxes in the bottom panel indicate samples used for *in vitro* transcription assays (Figs. 1 and S1). Both samples of derivatized H-NS were used for TEN-map experiments.

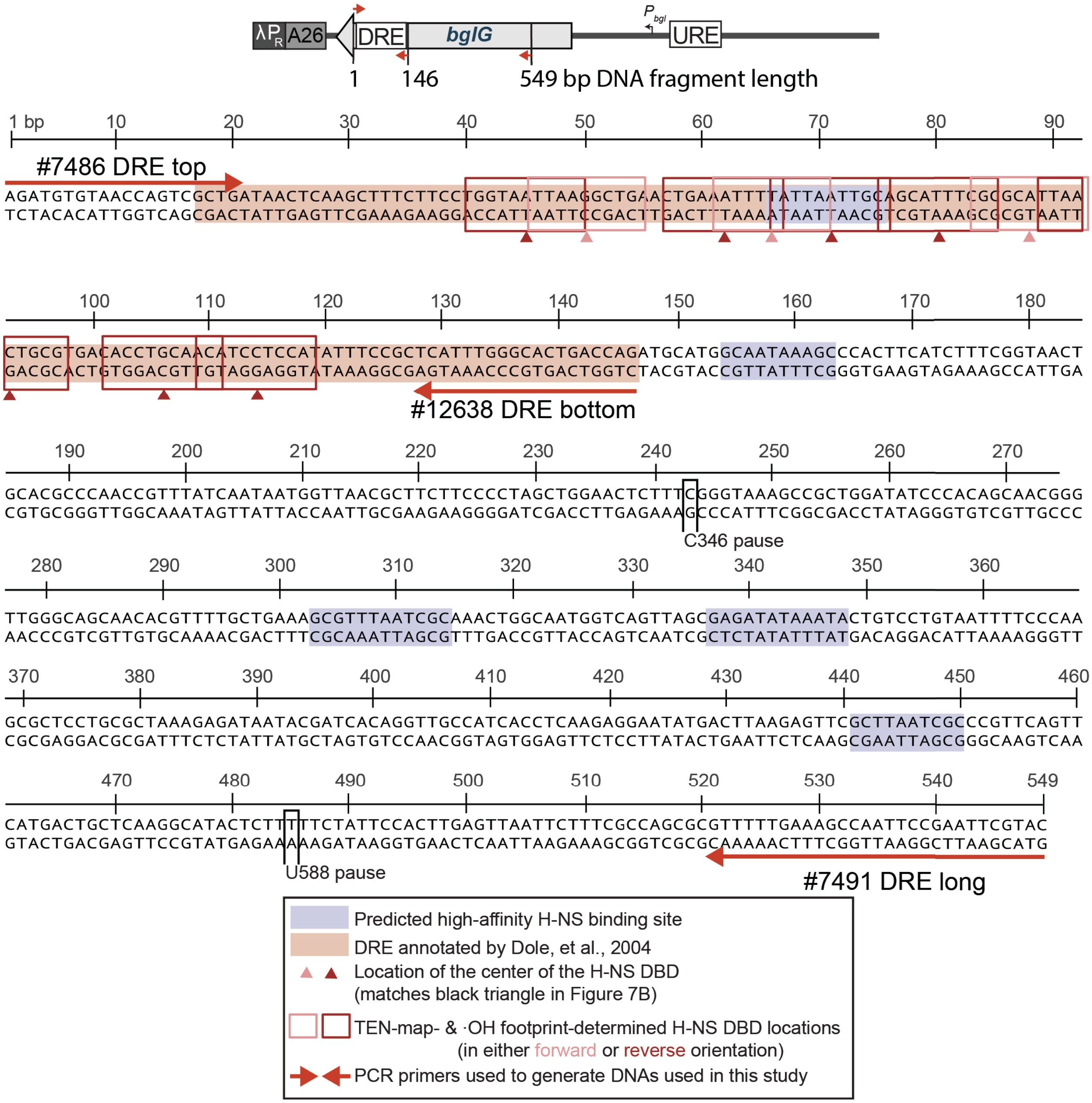


**Figure S3. DNA sequence of *bgl* DRE fragment used in TEN-map and ·OH footprinting experiments.**

Schematic (top) of the *in vitro* transcription template (see Figures 1 and S1) and the ends of the DNA fragments used in TEN-map and footprinting experiments (red arrows indicate PCR primer positions). The 146-bp DRE DNA fragment was generated by PCR using 5′ $^{32}$P-end-labeled primers to label either the top strand (primer #7486) or the bottom strand (primer #12638). The 549 bp DRE fragment was generated using primers #7486 and #7491 and labeled on the top strand. The DRE (Dole et al., 2004) is highlighted in light pink. High-affinity H-NS binding sites identified by Virtual Footprint (http://www.prodoric.de/vfp/vfp_regulon.php) are indicated by blue highlight. The sequence is numbered from the start of the 146 bp DRE fragment with the 346 and 588 pauses numbered based on transcript length (Figure 1) located at 243 and 486, respectively. The locations of H-NS DBDs in Figure 6C are indicated with light pink (forward) and dark red (reverse) open boxes and triangles at the binding site center (see black triangle in Figure 5).

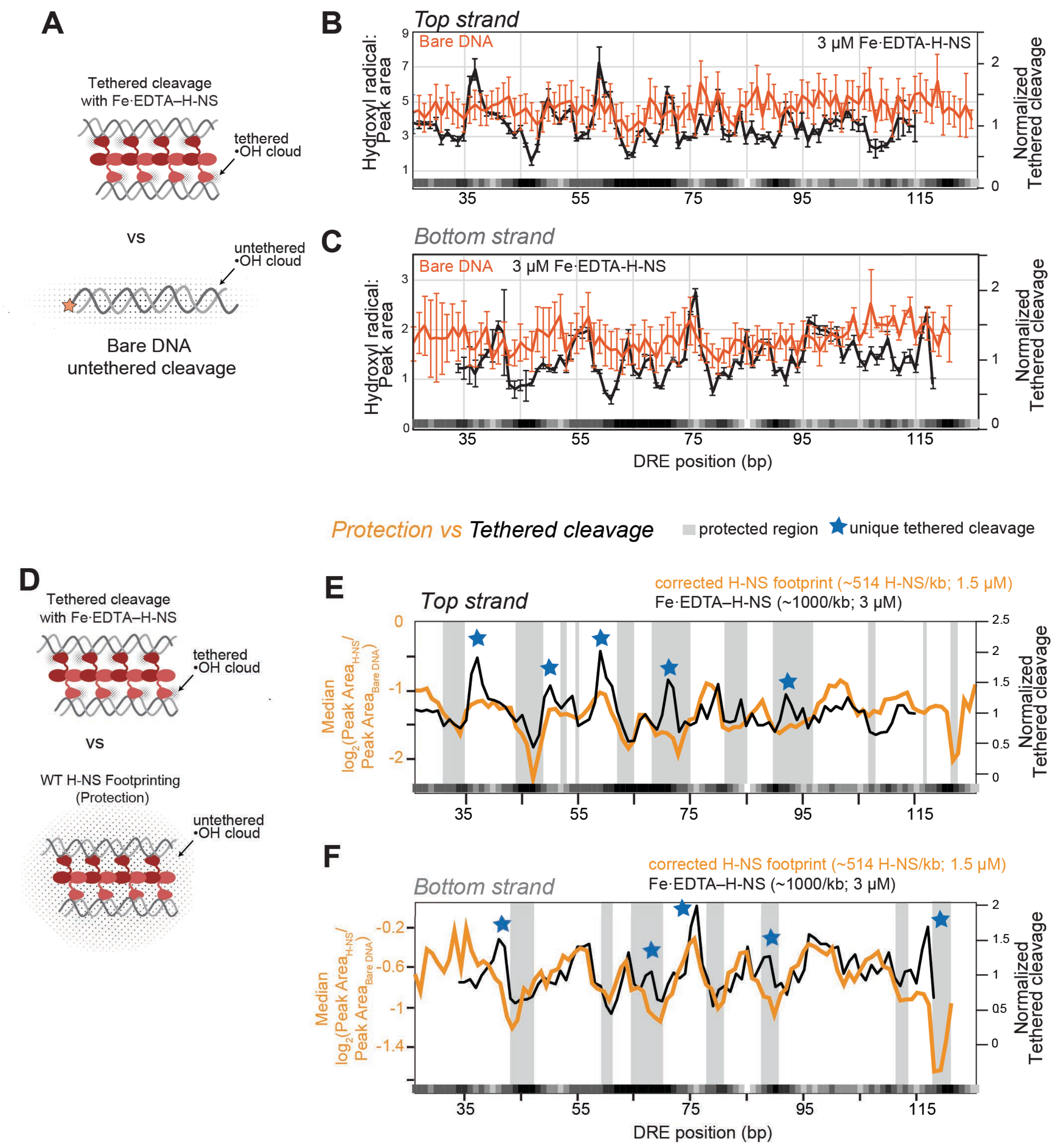


**Figure S4. TEN-map cleavage patterns differ from background DNA cleavage of H-NS footprints generated by ·OH from untethered $Fe^{2+}$–EDTA.**

(A) To verify that TEN-map cleavage reveals H-NS binding sites, the tethered cleavage pattern (top) was compared to ·OH cleavage of bare DRE DNA (bottom). Black dots represent the possible reach of ·OH cleavage in the assay. Peak area of ·OH cleavage of 146-bp DRE DNA top (B) or bottom (C) strand in the absence of H-NS shows intrinsic cleavage propensity of the DRE sequence. The intrinsic cleavage pattern of bare DNA (red) differed from the tethered cleavage pattern generated by 3 μM Fe·EDTA–H-NS (1027 H-NS/kb) on both strands (black traces). The difference between these lines indicates specific cleavage by $Fe^{2+}$·EDTA–H-NS. Error bars are standard deviation of three replicates.
(D) Comparison of the diffusion of ·OH in the TEN-map (top) and the ·OH footprint (protection; bottom)

experiments. Black dots represent the possible reach of ·OH cleavage in the assay.
(E) Orange line shows the ·OH footprints of wild-type H-NS on the top strand corrected by subtracting the bare DNA cleavage pattern from the pattern of ·OH cleavage of the bottom strand in the presence of wild-type H-NS (1.5 μM; Same trace as in Figure 4C). Protected regions (valleys in protection pattern) calculated from concentration-dependent slope estimates are indicated by shaded boxes (see Figure 4). Black line shows TEN-map cleavage by $Fe^{2+}$·EDTA–H-NS (3 μM; 1027 H-NS/kb) demonstrating its distinct pattern, including cuts within H-NS footprints. The most notable cuts attributable to tethered $Fe^{2+}$·EDTA are indicated by blue stars.
(F) Same as (E) but for the bottom strand.

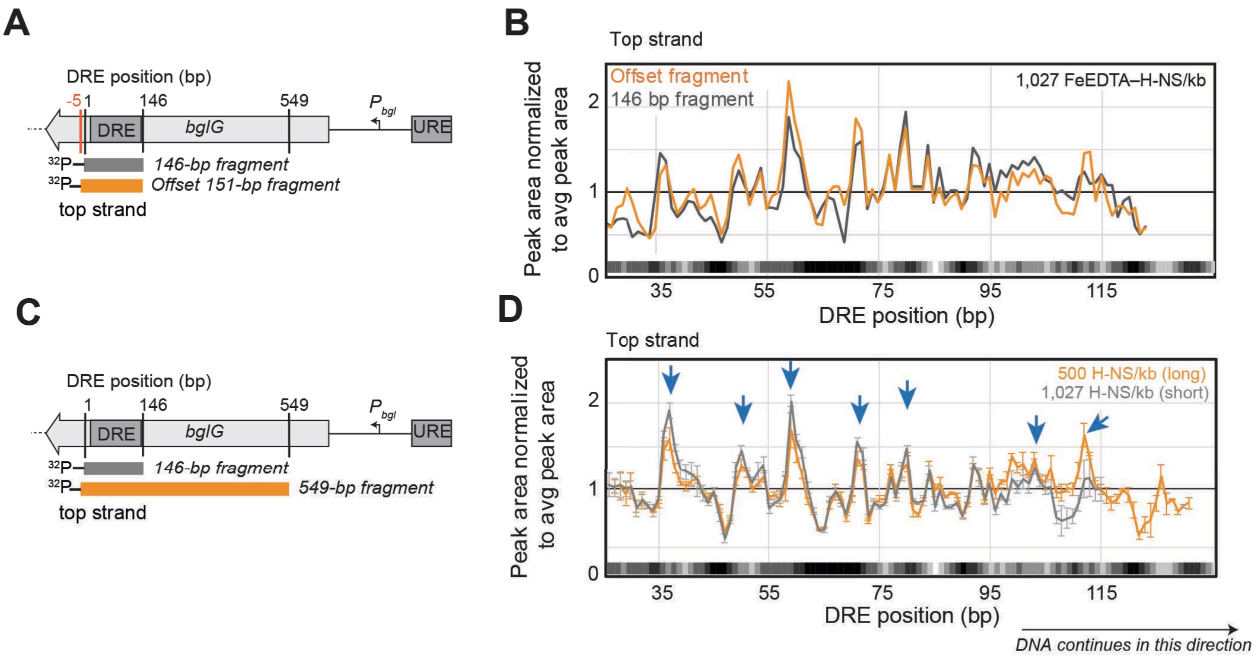


**Figure S5. TEN-map cleavage is unaffected by DNA ends or flanking DNA.**

(A) Schematic of the DNA templates used to test if H-NS binding is sequence-specific or initiates from the end of the DNA. An alternative PCR primer (orange line) was used to shift the DNA end of the top strand by 5 bp relative to the 146-bp fragment to create the offset fragment (orange box).
(B) TEN-map cleavage peak areas for the 146-bp fragment (black) and the 151-bp offset DNA fragment (orange). Cleavage and quantification were performed as described in Methods and show averages from two independent experiments.
(C) Comparison of the short (146 bp, gray box) and long (549 bp, orange box) DRE DNA fragments to test effects of DNA length on arrangement of H-NS DBD within a filament. Both fragments were labeled on the top strand.
(D) TEN-map cleavage patterns for standard (gray) or long (orange) DNAs. Gray line matches data in Figure 2C. To form filaments on long DNA, a mixture of 60% wild-type H-NS and 40% $Fe^{2+}$·EDTA–H-NS was used because the concentration of $Fe^{2+}$·EDTA–H-NS alone was too low to form pure $Fe^{2+}$·EDTA–H-NS filaments. Error bars show standard deviation of three replicates. Blue arrows indicate peaks of cleavage.

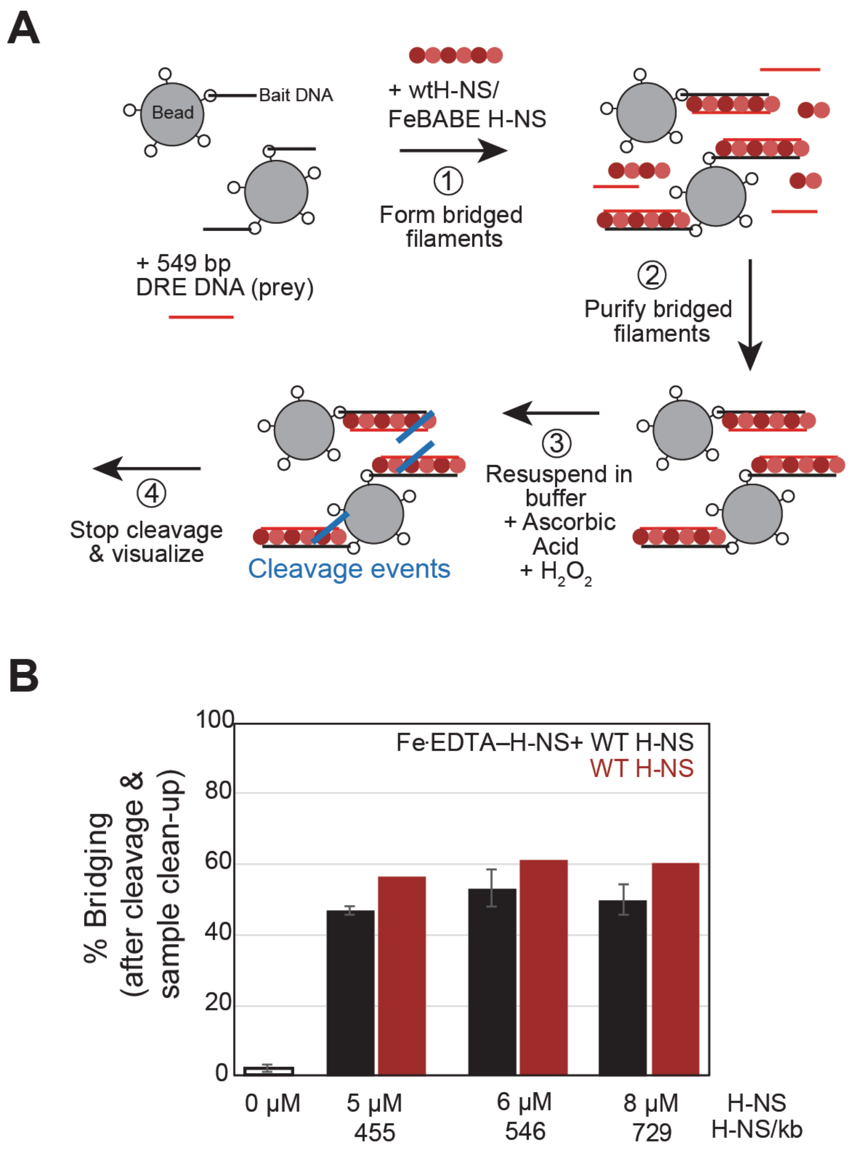


**Figure S6. TEN-map cleavage of obligately bridged filaments formed on beads.**

(A) Set-up of bridged cleavage assay. Filaments were formed with a 60-40 mix of wild-type H-NS to $Fe^{2+}$·EDTA–H-NS over a range of concentrations (0, 5, 6 and 8 μM H-NS or 0, 455, 546, and 729 H-NS/kb) in the presence of streptavidin paramagnetic beads containing immobilized 685-bp DNA (van der Valk et al., 2017). H-NS dimers (conjugated or wild-type) are shown as alternating dark and light red spheres to indicate head–head, tail–tail dimerization. The 549-bp DRE DNA (prey DNA) was not tethered to beads and was labeled on the top strand. Bridged filaments were recovered by magnetic separation and gently resuspended in 20 μL Filament Buffer (see Methods) followed by rapid addition of 1 μl 50 mM ascorbic acid and 4 μL 12.5 mM H2O2 to induce cleavage. Reactions were stopped with 22.4 mM thiourea, extracted with phenol, recovered by ethanol precipitation, and separated by denaturing 7% PAGE.

(B) The percentage of prey DNA recovered by bridging in the presence of either wild-type H-NS (red) or a mixture of wild-type H-NS and $Fe^{2+}$·EDTA–H-NS (black). Error bars show a standard deviation of three replicates, and bridging with WT H-NS was performed twice.

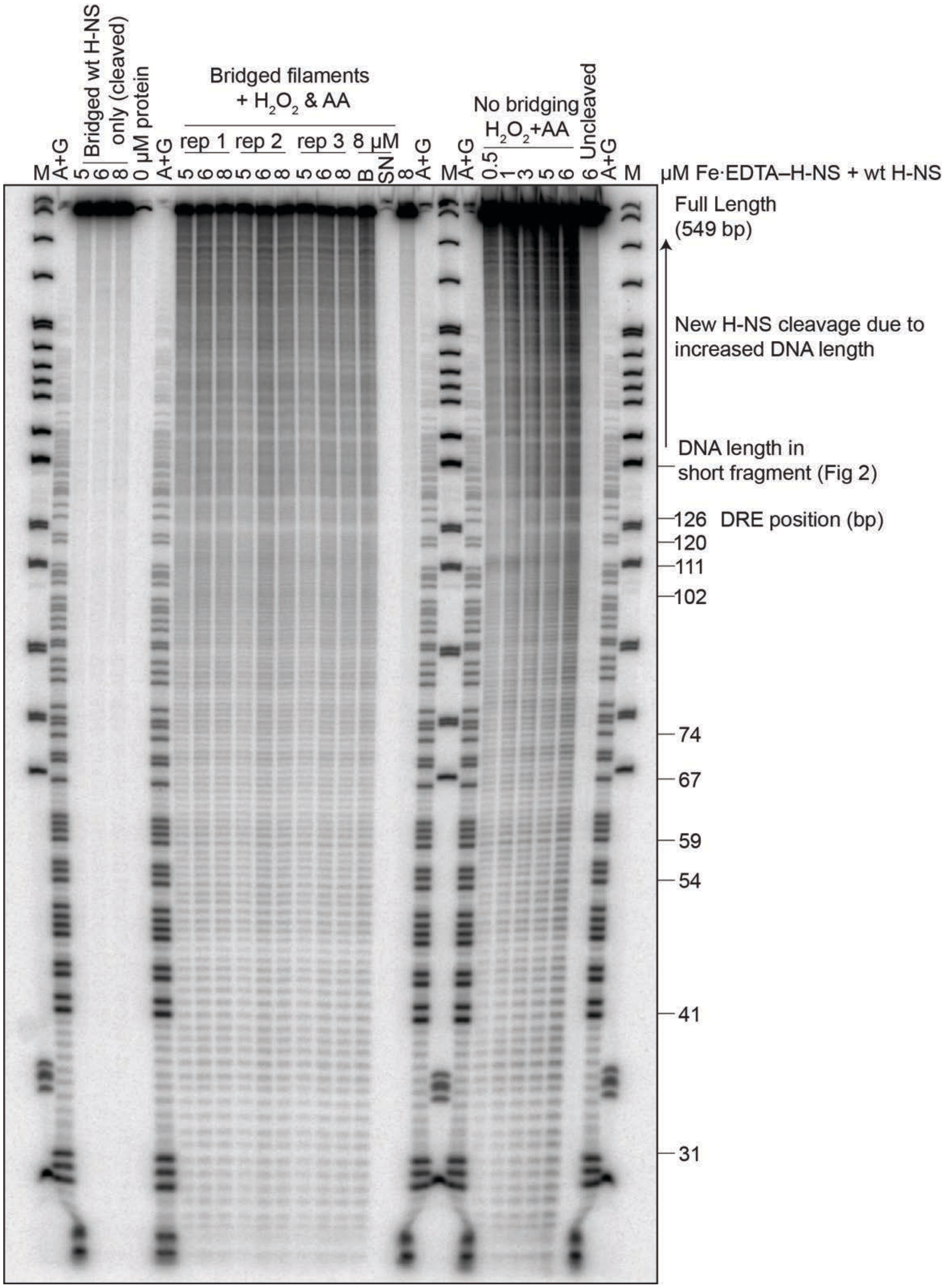


**Figure S7. TEN-map cleavage of obligately bridged DNA versus in solution.**

Denaturing 7% PAG showing cleavage products generated by Fe·EDTA–H-NS in filaments formed on beads (obligately bridged filaments) or in solution (no obligate bridging). H-NS used to form filaments was 40% Fe·EDTA–H-NS and 60% wild-type H-NS at the indicated concentrations. 8 μM H-NS lanes marked "B" and "SN" indicate filaments formed with 8 μM H-NS, cleaved, and then magnetically separated into supernatant (SN), which contained DNA that dissociated from the beads, and bound fractions ("B", obligately bridged). Non-obligately bridging filaments were formed on the 549-bp DRE DNA without beads or bait DNA present at indicated concentrations of H-NS. A+G, Maxam–Gilbert A and G sequencing ladder. M, MspI digest pBR322.

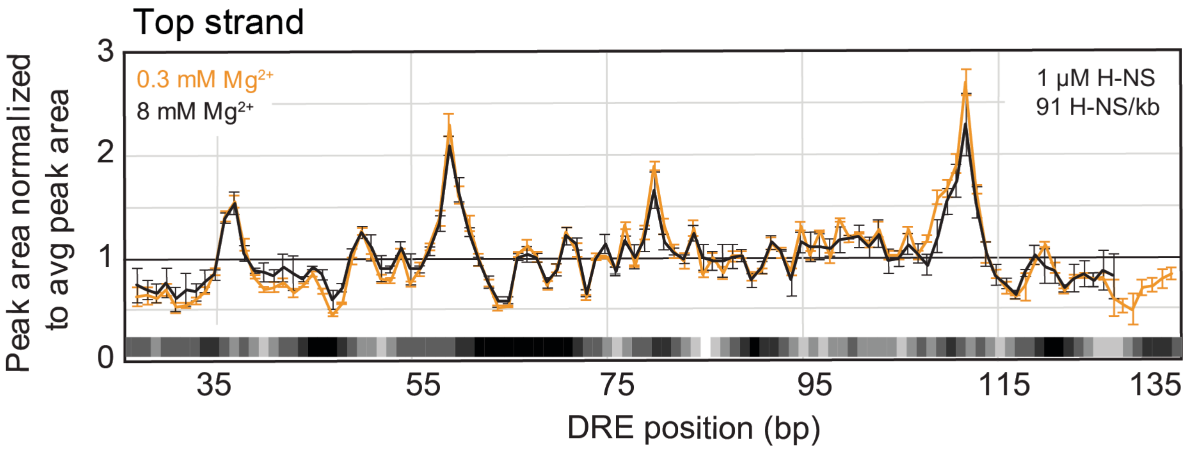


**Figure S8. H-NS DBDs bind DNA similarly at low and high $Mg^{2+}$.**

TEN-map cleavage pattern of the 549-bp DRE fragment ($^{32}P$-labeled top strand) at 1 μM H-NS (91 H-NS/kb) in the presence 8 mM $Mg^{2+}$ (black) or 0.3 mM $Mg^{2+}$ (orange). H-NS added was a 60-40 mixture of wild-type and $Fe^{2+}$·EDTA–H-NS to match experiments done on beads (Figure 3A). Error bars represent the standard error of at least three replicates.

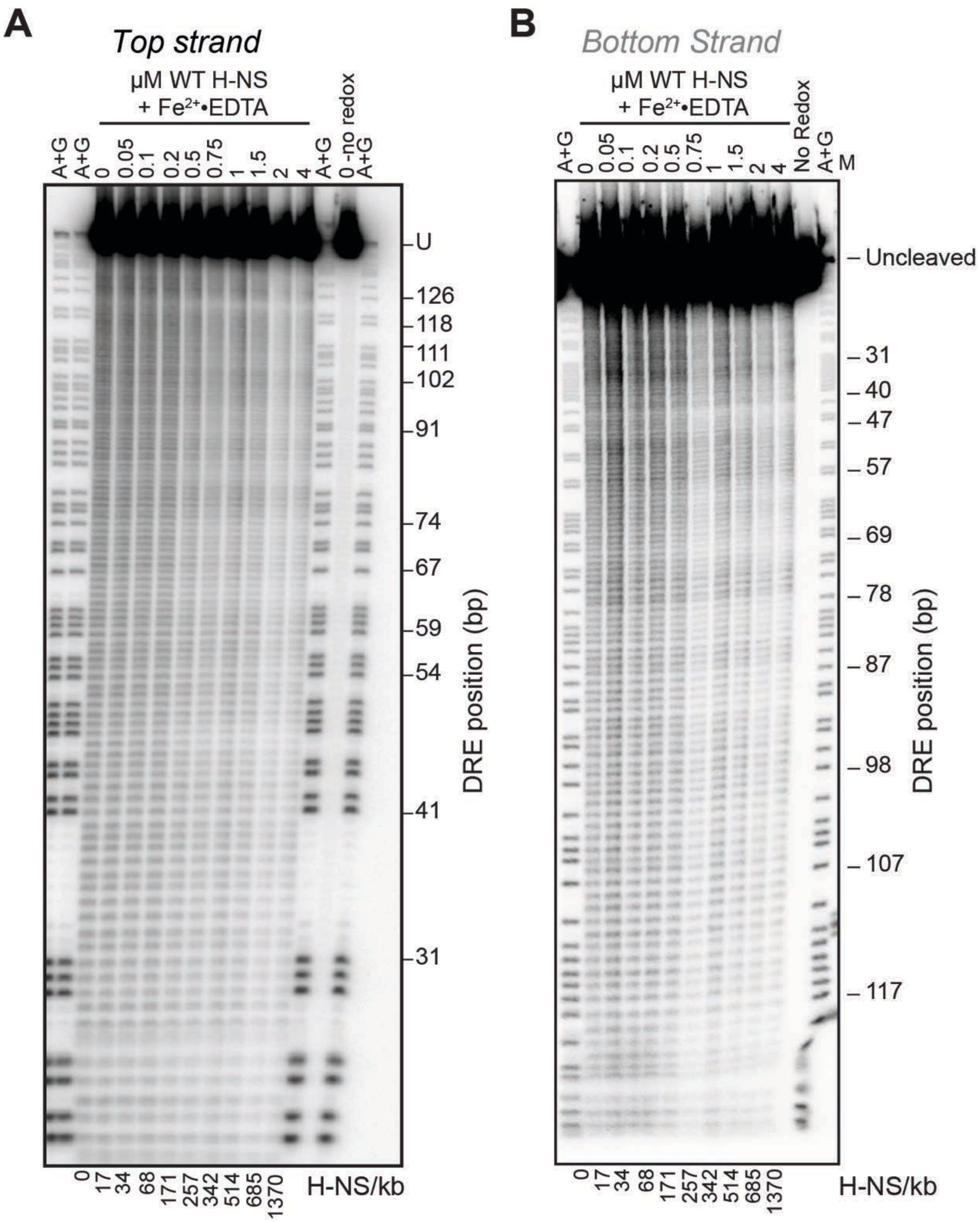


**Figure S9. WT H-NS ·OH footprinting reveals sites of H-NS protection on the DRE DNA in bridged and linear filaments.**

Denaturing 7% PAG showing ·OH cleavage pattern of 146-bp DRE DNA $^{32}P$-labeled on either the on the top (A) or bottom (B) strand in the presence or absence of wild-type H-NS filaments formed at 0.05, 0.1, 0.2, 0.5, 0.75, 1, 1.5, 2, or 4 μM H-NS (17, 34, 68, 171, 251, 342, 514, 685, 1370 H-NS/kb DNA). The ratio of H-NS/kb is indicated below the gel. Wild-type H-NS plus untethered $Fe^{2+}$·EDTA footprinting experiments were performed by sequential addition of $H_2O_2$, ascorbic acid, and free $Fe^{2+}$·EDTA. Gels are representative of at least 3 experiments.

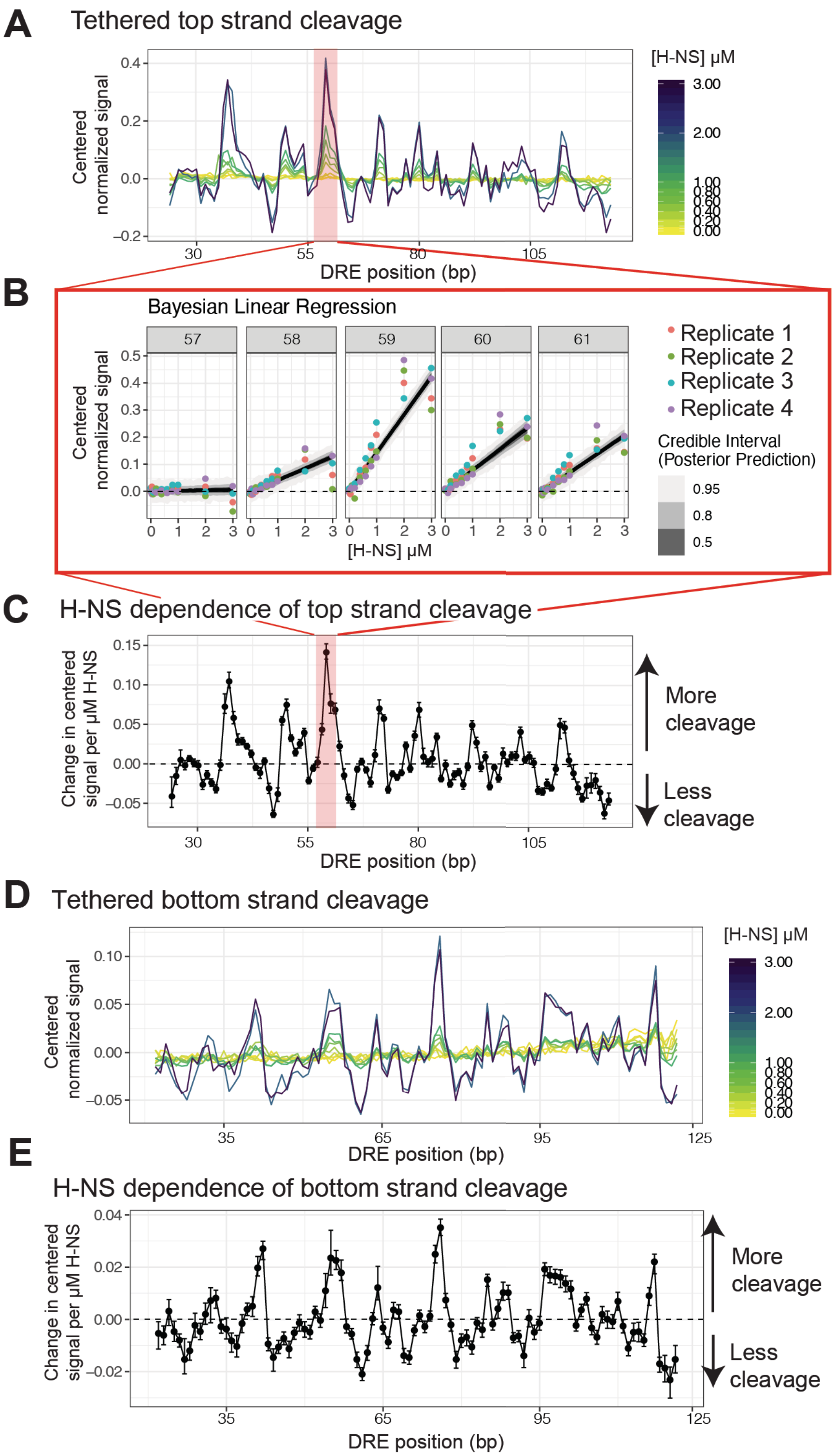


**Figure S10. Quantification of TEN-map cleavage by linear regression model.**
(A) Cleavage of the top strand of the DRE DNA fragment as a function of $Fe^{2+}$·EDTA–H-NS concentration (yellow-to-purple gradient). Cleavage signal were corrected for total signal in each lane and then normalized to the average cleavage signal and centered on the mean.
(B) Example of slope determination by linear regression for positions 57-61 on the top strand. The

relationship between mean-centered signals and the $Fe^{2+}$·EDTA–H-NS concentration (from 0 to 3 μM) on the bottom strand was determined for each bp by using a Bayesian linear regression model.
(C) H-NS dependence of cleavage of the bottom strand as determined from Bayesian linear regression model. Error bars indicate credible intervals on the slope estimates.
(D) Same as B, but for the bottom strand.
(E) Same as D, but for the bottom strand.

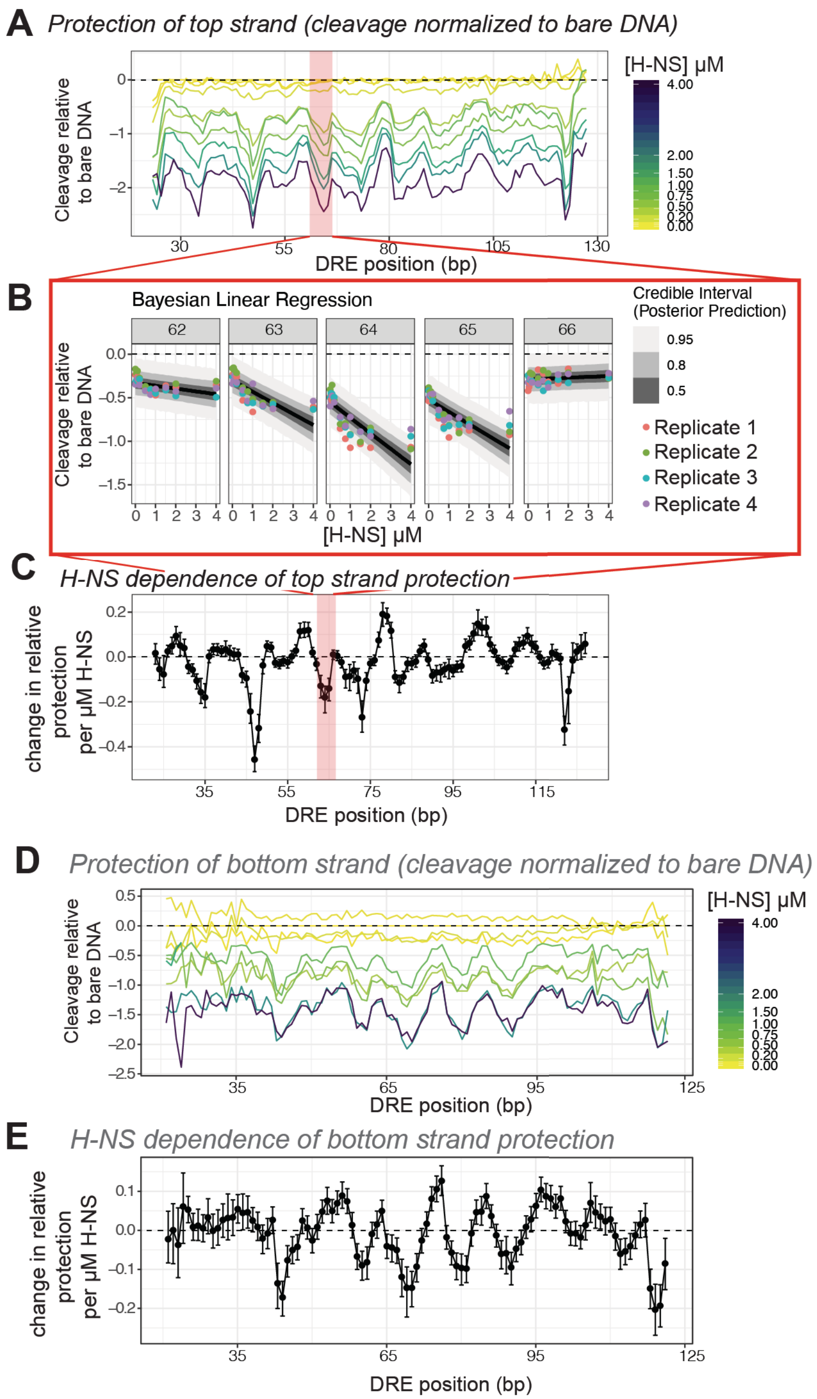


**Figure S11. Quantification of ·OH footprinting of H-NS filaments using a linear regression model.**
(A) Protection of the top strand of the DRE DNA fragment from ·OH cleavage as a function of H-NS concentration (yellow-to-purple gradient). (Same as Figure 4B). The ·OH footprinting pattern for H-NS was quantified by calculating the ratio of signal at each position compared to cleavage of bare DNA. The median of 4 replicates was calculated, but error bars are removed for clarity.
(B) Example of slope determination by linear regression for the for positions 62-66 on the top strand. The change in cleavage as a function of H-NS concentration (slope) was fit to a Bayesian linear regression model.
(C) Changes in protection relative to bare DNA (slopes from linear regression fits) for the top strand.

Error bars indicate 95% credible intervals for the slope estimates.
(D) Same as A, but for the bottom strand. Data also shown in Figure 4C.
(E) Same as C, but for the bottom strand.

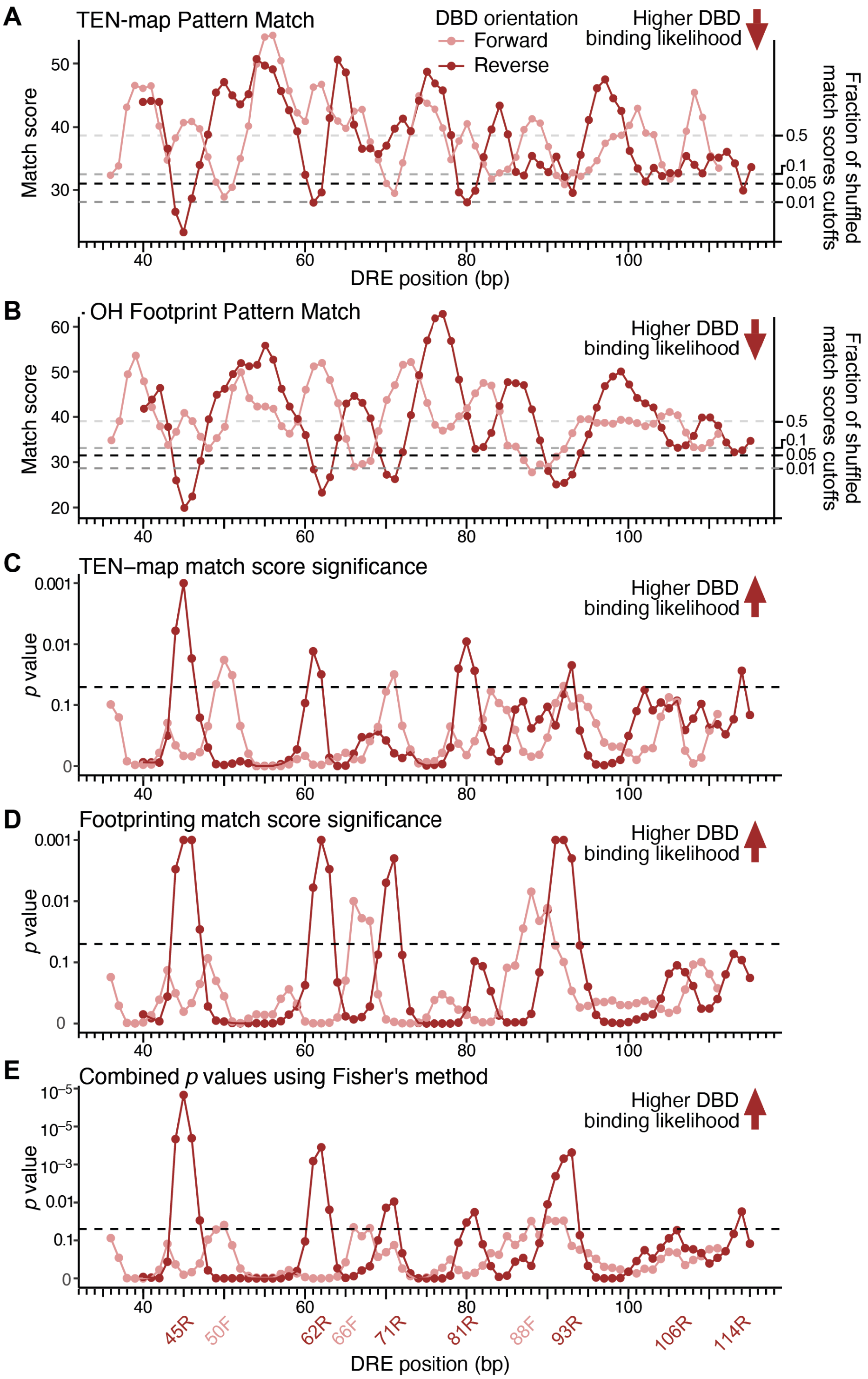


**Figure S12. DBD location probabilities from comparison of molecular modeling–based predictions to TEN-map and ·OH footprint results.**

(A) Match scores based on Manhattan distances between predicted and observed cleavage signals for TEN-map analysis of the *bgl* DRE fragment (see Figure 5). A lower match score indicates a closer match of the data to the prediction for a given location and orientation (light red, forward; dark red, reverse). The fraction of matches to randomized data that give match scores equal or less than the indicated value

in a Monte Carlo permutation test is shown by the dotted lines and righthand y-axis (see Methods).
(B) Match scores for ·OH footprint data as described for panel A.
(C) Monte Carlo permutation test *p* values (see Methods for details) for DBD locations based on TEN-map data derived from the analysis shown in panel A. A lower *p* value indicates a lower probability that the agreement between the observed and predicted cleavage signals had arisen by chance for a given location and orientation (light red, forward; dark red, reverse)..
(D) Monte Carlo permutation test *p* values for DBD locations based on ·OH footprint data as described for panel C.
(E) *p* values calculated for the combined TEN-map and ·OH data (see Methods).
